# Conformation and dynamics of the kinase domain drive subcellular location and activation of LRRK2

**DOI:** 10.1101/2020.07.13.198069

**Authors:** Sven H. Schmidt, Jui-Hung Weng, Phillip C. Aoto, Daniela Boassa, Sebastian Mathea, Steven Silletti, Junru Hu, Maximilian Wallbott, Elizabeth A Komives, Stefan Knapp, Friedrich W. Herberg, Susan S. Taylor

## Abstract

In a multi-tiered approach, we explored how Parkinson’s Disease-related mutations hijack the finely tuned activation process of Leucine-Rich Repeat Kinase 2 (LRRK2) using a construct containing the ROC, Cor, Kinase and WD40 domains (LRRK2_RCKW_). We hypothesized that the N-terminal domains shield the catalytic domains in an inactive state. PD mutations, type-I LRRK2 inhibitors, or physiological Rab GTPases can unleash the catalytic domains while the active kinase conformation, but not kinase activity, is essential for docking onto microtubules. Mapping solvent accessible regions of LRRK2_RCKW_ employing hydrogen-deuterium exchange mass spectrometry (HDX-MS) revealed how inhibitor binding is sensed by the entire protein. Molecular Dynamics simulations of the kinase domain elucidated differences in conformational dynamics between wt and mutants of the DYGψ motif. While all domains contribute to regulating kinase activity and spatial distribution, the kinase domain, driven by the DYGψ motif, coordinates domain crosstalk and serves as an intrinsic hub for LRRK2 regulation.

## Introduction

Although Parkinson’s Disease (PD), first described in 1817 (1), is the second most common neurodegenerative disease to date, there are still no cures available (2–4). A primary reason for this lack of progress is that the underlying molecular processes that drive PD are still barley understood. Leucine rich repeat kinase 2 (LRRK2) is a good target to improve our knowledge of these underlying processes as mutations within this protein are the most common cause for genetically driven forms of PD (5, 6). LRRK2 is a complex and multifunctional protein consisting of seven closely interacting structural domains (7, 8). LRRK2 belongs to the protein family of ROCO proteins, which are defined by containing a GTPase domain called Ras of Complex (Roc) followed by a COR domain in combination with other domains. A special feature of LRRK2, as well as some but not all of the ROC:COR proteins, is that it contains both, a kinase domain and a GTPase domain so that two catalytically active core domains are embedded within the same polypeptide chain (9–11).

Familial PD mutations in LRRK2 lead to altered cellular phenotypes such as microtubule (MT) associated filament formation, impeded vesicular trafficking, as well as changes in nuclear morphology (12–19). Some of these PD mutations are found in the kinase domain, for example G2019S leads to increased kinase activity, and so kinase activity is thought to be a major component of LRRK2 regulation. Yet, some potent kinase inhibitors such as type-I inhibitor MLi-2 (Merck LRRK2 Inhibitor 2) as well as other kinase mutations such as I2020T with unchanged or reduced catalytic activity also lead to the same disease-like cellular phenotypes (17, 20, 21). Thus, it is clear that the kinase domain plays an important pathogenic role but that its activity alone is insufficient to describe the complexities of LRRK2 regulation. With five scaffolding and two catalytically active domains the interplay and how these domains effect and regulate each other is very complex and allows for multilayered intrinsic cross-regulation between the kinase and the GTPase domains. However, due to the lack of high-resolution structural data not much was known about these interactions and how they intrinsically regulate LRRK2. LRRK2-dimerization, and interactions with other proteins like Rab and 14-3-3 proteins further increase the complexity of LRRK2 regulation (22–33).

To unravel the complexity of LRRK2 regulation we used a multi-tiered approach beginning with a cell-based assay for filament formation, a process that correlates with LRRK2 docking onto microtubules (MTs) (14). Specifically, we used live cell imaging to examine the spatial and temporal distribution of full length wt and G2019S LRRK2 in HEK293T cells in the presence and absence of LRRK2 kinase inhibitors. To explore the regulatory functions that are embedded in the catalytically inert N-terminal domains (NTDs) of LRRK2 and to discriminate between the catalytic C-terminal domains (CTDs) we engineered a construct, LRRK2_RCKW_, that contains only the CTDs (<u>R</u>OC, <u>C</u>OR, <u>K</u>inase and <u>W</u>D40 domains).

This construct contains both catalytic moieties of LRRK2 and exhibits functional kinase and GTPase activity embedded in the same protein. It was shown previously that kinase activity of LRRK2 is dependent on the presence of the ROC:COR domain and the C-terminus (34, 35). On the protein level, we investigated the kinase activity of wt LRRK2_RCKW_ and several variants using LRRKtide and Rab8a as substrates. We focused in particular on the DYGψ motif in the kinase domain which is a hot spot for PD mutations and where we showed previously that the tyrosine (Y2018), conserved as a Phe in most other protein kinases, is a critical part of the switch mechanism that leads to LRRK2 activation (36). Using hydrogen deuterium exchange mass spectrometry (HDXMS) we then zoomed in to the level of LRRK2 peptides and demonstrated that LRRK2_RCKW_ is a well-folded protein. We continued to map changes in the solvent exposure of the LRRK2_RCKW_ domains following binding of MLi-2 to the kinase domain, which provided an allosteric portrait of the kinase domain as a hub for driving long-range conformational changes. At yet another level, we used Gaussian molecular dynamics (GaMD) simulations as a computational microscope to observe at an atomistic level how single amino acid mutations in the kinase domain contribute to the intrinsic dynamic regulation of LRRK2.

Our multi-scale approach allowed us to achieve a better understanding of the intrinsic regulation processes of LRRK2 and emphasized the crucial role of the DYGψ motif in regulating LRRK2 structure and function. We hypothesized that the N-terminus of LRRK2 (ARM:ANK:LRR) plays a regulatory role acting as a “lid”, which can be “unleashed” from the catalytic (RCKW) domains by kinase inhibitors or constitutively by some of the PD mutations. In addition to shielding the catalytic domains, the N-terminal lid mediates interactions with other proteins that contribute to activation and subcellular localization (26–29, 37, 38). Strikingly, the resulting LRRK2 model for activation and subcellular localization closely resembles the activation process of Raf kinases, thereby underlining the plausibility of our model and the central concept of kinases serving as the hub for driving conformational switching in multi-domain signaling proteins.

## Methods

### HEK293T Cell culture and Transfection

For expression of each Flag-Strep-Strep-tagged (FSS) LRRK2_RCKW_ construct cells on ten 15 cm Ø cell culture dishes were transfected. Therefore, 1.0×10^7^ HEK293T cells (human embryonic kidney cells carrying the temp sensitive mutant of the SV-40 large T-antigen, *DSMZ*, DSMZ-No:ACC635) were seeded per dish and incubated for 24 h at 37°C and 5% CO_2_ in Dulbecco’s Modified Eagle Medium (DMEM) high glucose (w. L-Glutamine; w.o. Sodium Pyruvate, *biowest*) supplemented with 10% fetal bovine serum (FBS). In the following, transfections were performed by adding a 30 min preincubated mixture of 15 μg plasmid DNA (pCDNA3.0-FSS-LRRK2_RCKW_ (aa1327-2527); NM_198578), 150 μl PEI (Polyethylenimine) (1 μg/μl) and 1.5 ml DMEM high glucose per dish. Medium was exchanged with fresh DMEM (high glucose, +10% FBS) after 24 h. After another 24 h cells were harvested and stored at −20°C before use.

### LRRK2_RCKW_ Transfection and Expression in Sf9 cells

Exponentially growing TriEx Sf9 insect cells (cells are derived from a high-yielding clone of *Spodoptera frugiperda* cell IPLB Sf21-AE (ATCC CRL-1711), *Novagen*, Prod.-No.: 71023-3) were diluted to a density of 2×10^7^ cells/mL. To initiate LRRK2_RCKW_ expression, a high-titer virus suspension was added in the ratio 1:64. The viruses had been generated using the Bac-to-Bac expression system (*Invitrogen*) and the expression vector pFB-6HZB. Accordingly, the expressed protein was composed of an N-terminal His_6_-Z tag and the LRRK2 residues 1327 to 2527. The infected cells were incubated in 800 mL aliquots in shaker flasks (66 h, gentle agitation, 27°C), harvested by centrifugation and stored at −20°C.

### Purification of overexpressed Flag-Strep-Strep-tagged LRRK2_RCKW_ constructs from HEK293T Cells

Each pellet was resuspended in 10 ml of fresh ice cold lysis buffer (25 mM Tris-HCl pH7.5, 150 mM NaCl, 10 mM MgCl_2_, 0.5% Tween 20, 500 μM GTP, cOmplete™ EDTA free protease inhibitor cocktail [*Roche*], PhosSTOP™ [*Roche*]) and incubated for 30 min at 4°C on a rotating wheel to lyse cells. In the following, cell debris were removed by centrifugation at 42,000 xg and 4°C for 40 min and a filtration step (0.45 μm sterile filter). The supernatant was loaded onto a Streptactin Superflow column (0.5 mL bed volume, *IBA Goettingen*) and purification was performed according to the manufacturer’s protocol while all buffers were additionally supplemented with 500 μM GTP (*Biolog Life Science Institute*) and 10 mM MgCl_2_. The purified LRRK2_RCKW_ constructs were stored at −80°C containing 10% Glycerol and 0.5 mM TCEP. The LRRK2_RCKW_ construct concentrations were determined after Bradford (39).

### Purification of overexpressed LRRK2_RCKW_ constructs from Sf9 cells

The LRRK2_RCKW_ expression construct contained an N-terminal His_6_-Ztag and a TEV protease cleavage site. For purification, the Sf9 cell pellets were washed with PBS, resuspended in lysis buffer (50 mM HEPES pH 7.4, 500 mM NaCl, 20 mM imidazole, 0.5 mM TCEP, 5% glycerol, 5 mM MgCl_2_, 20 μM GDP) and lysed by sonication. The lysate was cleared by centrifugation and loaded onto a Ni NTA column. After vigorous rinsing with lysis buffer the His_6_-Ztagged protein was eluted in lysis buffer containing 300 mM imidazole. Immediately thereafter, the eluate was diluted with a buffer containing no NaCl, in order to reduce the NaCl-concentration to 250 mM and loaded onto an SP sepharose column. His_6_-ZTEV-LRRK2_RCKW_ was eluted with a 250 mM to 2.5 M NaCl gradient and treated with TEV protease overnight to cleave the His_6_-Ztag. Contaminating proteins, the cleaved tag, uncleaved His_6_ZTEV-LRRK2_RCKW_ and TEV protease were removed in another combined SP sepharose Ni NTA step. Finally, LRRK2_RCKW_ was concentrated and subjected to gel filtration in storage buffer (20 mM HEPES pH 7.4, 800 mM NaCl, 0.5 mM TCEP, 5% glycerol, 2.5 mM MgCl_2_, 20 μM GDP) using an AKTA Xpress system combined with an S200 gel filtration column.

### Live Cell Imaging

Time-lapse imaging was conducted using an Olympus FluoView1000 laser scanning confocal microscope equipped with a CO_2_ and temperature-controlled chamber (at 37°C) and a 60X 1.42 NA objective lens. The HEK293T cells (*CCLV*, RRID:CVCL_0063) were imaged in HBSS supplemented with 5% fetal bovine serum and 10 mM HEPES and maintained at 37°C throughout the experiment. YFP fluorescence and the differential interference contrast were collected, 515 nm excitation/530–630 nm emission. Images were recorded every 5 to 11 min for a total duration of 2 h or 14 h as specified in figure legends, sampling speed 2 μs/pixel, image size 640×640 pixels in 15-20 z slices/area (step size 1 μm). Cells were imaged before and after treatment with inhibitors MLi-2 (100 nM) or rebastinib (100 nM). For washout experiments, cells were washed 5 times with imaging media, 2 min each, and then the imaging session was resumed using the same settings as before washout. Image processing such as 3D volume rendering and 4D movie generation was done using the Imaris Software (Bitplane AG, St. Paul, MN) and the Image J software (rsb.info.nih.gov/ij).

### Immunofluorescence and laser confocal imaging

HEK293T cells were seeded onto 6-well dishes containing poly-D-lysine-coated glass coverslips or onto 35mm poly-D-lysine-coated glass bottom dishes (MatTek Corporation, Ashland, MA, USA). For HEK293T cell transfection, 1 μg of Flag-Strep-Strep-(FSS)-tagged LRRK2_RCKW_ cDNA and Lipofectamine 2000 reagent (ThermoFisher Scientific, USA) were used according to the manufacturer’s protocol. After incubation for 48 h at 37 °C cells were treated for 2 h with the LRRK2 inhibitor MLi-2. Subsequently cells were fixed with 4% paraformaldehyde in phosphate-buffered saline (PBS) for 15 minutes at room temperature. Cells were washed in PBS, permeabilized in 0.1% Triton X-100, and blocked in 1% BSA, 50 mM glycine and 2% normal donkey serum. A rabbit anti-Flag antibody (Abnova Company, Cat. No. PAB0900) was mixed 1:200 in blocking buffer diluted five-fold in PBS. The primary antibody solution was applied to the cells for 1 h at room temperature. The secondary antibody (donkey-anti-rabbit-Alexa568, Invitrogen, Cat. No. A10042) was also diluted (1:100) in the blocking buffer diluted five-fold in PBS and applied for 1 h at room temperature. Samples were mounted with the antifade agent ProLong Gold with DAPI (ThermoFisher Scientific, USA). The Olympus Fluoview 1000 laser scanning confocal microscope utilizing a 60X oil immersion objective lens with a numerical aperture of 1.42 was used for confocal imaging. Z-stack images were acquired with a step size of 0.3 microns and processed using the Fiji software package (40). Cells expressing the different mutants were assessed for the presence of clear filamentous structures and quantified in two independent experiments in case of the untreated cells and in one experiment for the MLi-2 treated cells.

### Rab8a phosphorylation by LRRK2_RCKW_ variants and pathogenic mutations

Phosphorylation of T72 of Rab8a was measured via Western Blotting using a pT72 specific antibody. Prior to the blotting step onto a nitrocellulose membrane an *in vitro* kinase assay using kinase buffer (25 mM TRIS/HCl, pH 7.5, 50 mM NaCl, 10 mM MgCl_2_, 1 mM ATP, 0.5 mM GTP, 0.1 mg/ml BSA and 1 mM DTT) and an SDS-PAGE were performed. For the kinase assay 2.5 μM (6xHis)-Rab8a (aa 6-175) were used as substrate for 200 nM of the LRRK2_RCKW_ variants. Rab8a was phosphorylated at 30 °C for 7 min at 650 rpm on a shaker. To stop the reaction 1xNu-PAGE LDS sample buffer (*Invitrogen*, Cat. No. NP0007) supplemented with 250 μM DTT was added followed by an incubation at 80 °C for 5-10 min. After SDS-PAGE and western blotting, membranes were blocked with 5% (w/V) BSA in TBS-T (1x Tris-buffered saline supplemented with 0.1% Tween20) for 1 h. Subsequently they were incubated overnight at 4 °C with the primary antibodies against pT72 (MJF-R20, *abcam*, Cat. No. ab231706) and the His-tag of Rab8a (anti-His-Antibody, *GE Healthcare, mouse*). Both were diluted (1:1000) in blocking buffer. Membranes were then washed three times with TBS-T. After secondary antibody incubation (anti-rabbit IRDye800 and anti-mouse IRDye680, 1:15000, *LiCOR*) for 1h at RT signals were detected using the Odyssey FC imaging system (*LiCOR*).

### Microfluidic Mobility Shift Kinase Assay (MMSKA)

To quantify the kinase activities of the LRRK2_RCKW_ variants MMSKA were performed using 1 mM ATP and 1 mM LRRKtide (RLGRDKYKTLRQIRQ-amide, *GeneCust*) as substrates. For these assays two stock solutions were prepared: a 2x concentrated (conc.) LRRKtide solution (1900 μM LRRKtide, 100 μM Fluorescein-LRRKtide, 2 mM ATP) and a 2x conc. LRRK2_RCKW_ variant solution (100-200 nM LRRK2_RCKW_ variant, 20 mM MgCl_2_, 1 mM GTP). All stock solutions were prepared using kinase buffer (25 mM TRIS/HCl, pH 7.5, 50 mM NaCl, 0.1 mg/ml BSA and 1 mM DTT). The reactions were started by mixing both solutions in a 1:1 ratio in 384 well plates. Reactions were performed at 30 °C and monitored for 60-90 min using a LabChip EZ Reader (*PerkinElmer*). The slope (conversion rate, [m]=%/min) of the percental conversion plotted against the time was determined using a linear fit model of Graph Pad Prism 6 and was converted into a reaction velocity ([v_0_]=μmol/min). For every construct at least three independent measurements were performed.

Titration assays using the high-affinity inhibitor MLi-2 (Merck, USA) were performed to determine the active protein concentrations of the LRRK2_RCKW_ variants. Therefore, 24 μL of Buffer A (25 mM TRIS/HCl, 50 mM NaCl, 20 mM MgCl_2_, 1 mM GTP, 1 mM DTT, 0.5 mg/ml BSA, 52.1/104.2 nM LRRK2_RCKW_ variant) were mixed with 1 μL of an MLi-2 dilution series (50x concentrated) prepared in 100% DMSO. To start the reaction 10 μL of this reaction mix was added to 10 μL of Buffer B (25 mM TRIS/HCl, 50 mM NaCl, 1 mM DTT, 0.5 mg/ml BSA, 1900 μM LRRKtide, 100 μM Fluorescein-LRRKtide, 360 μM ATP). The resulting conversion rates were plotted against the respective MLi-2 concentrations and to obtain the active protein concentrations the x-axes intersection of the respective linear fit was determined using Graph Pad Prism 6 (assuming a 1:1 binding of MLi-2).

### Hydrogen-deuterium exchange mass spectrometry

Hydrogen/deuterium exchange mass spectrometry (HDXMS) was performed using a Waters Synapt G2Si equipped with nanoACQUITY UPLC system with H/DX technology and a LEAP autosampler. The sample concentration was 5 μM in LRRK2 buffer containing: 20 mM HEPES/NaOH pH 7.4, 800 mM NaCl, 0.5 mM TCEP, 5% Glycerol, 2.5 mM MgCl_2_ and 20 μM GDP. The deuterium uptake was measured in LRRK2 buffer in the presence and absence of the kinase inhibitor MLi-2 (50 μM). For each deuteration time, 4 μL complex was equilibrated to 25 °C for 5 min and then mixed with 56 μL D_2_O LRRK2 buffer for 0, 0.5, 1 or 2 min. The exchange was quenched with an equal volume of quench solution (3 M guanidine, 0.1% formic acid, pH 2.66). The quenched sample (50 μL) was injected into the sample loop, followed by digestion on an in-line pepsin column (immobilized pepsin, Pierce, Inc.) at 15 °C. The resulting peptides were captured on a BEH C18 Vanguard pre-column, separated by analytical chromatography (Acquity UPLC BEH C18, 1.7 μM, 1.0 × 50 mm, Waters Corporation) using a 7-85% acetonitrile gradient in 0.1% formic acid over 7.5 min, and electrosprayed into the Waters SYNAPT G2Si quadrupole time-of-flight mass spectrometer. The mass spectrometer was set to collect data in the Mobility, ESI+ mode; mass acquisition range of 200–2,000 (m/z); scan time 0.4 s. Continuous lock mass correction was accomplished with infusion of leu-enkephalin (m/z = 556.277) every 30 s (mass accuracy of 1 ppm for calibration standard). For peptide identification, the mass spectrometer was set to collect data in MS^E^, ESI+ mode instead.

The peptides were identified from triplicate MS^E^ analyses of 10 μM LRRK2_RCKW_, and data were analyzed using PLGS 3.0 (Waters Corporation). Peptide masses were identified using a minimum number of 250 ion counts for low energy peptides and 50 ion counts for their fragment ions. The peptides identified in PLGS were then analyzed in DynamX 3.0 (Waters Corporation) using a cut-off score of 6.5, error tolerance of 5 ppm and requiring that the peptide be present in at least 2 of the 3 identification runs. The peptides reported on the coverage maps are those from which data were obtained. The relative deuterium uptake for each peptide was calculated by comparing the centroids of the mass envelopes of the deuterated samples vs. the undeuterated controls (41). For all HDX-MS data, at least 2 biological replicates were analyzed each with 3 technical replicates. Data are represented as mean values +/− SEM of 3 technical replicates due to processing software limitations, however the LEAP robot provides highly reproducible data for biological replicates. The deuterium uptake was corrected for back-exchange using a global back exchange correction factor (typically 25%) determined from the average percent exchange measured in disordered termini of various proteins (42). Deuterium uptake plots were generated in DECA (github.com/komiveslab/DECA) and the data are fitted with an exponential curve for ease of viewing (43).

### Gaussian accelerated Molecular Dynamics (GaMD) simulation

The LRRK2 kinase domain constructs for simulations were prepared using Phyre2 (44) with the crystal structure of Src kinase (PDBID: 1Y57) serving as an initial template. The model was separately then mutated to Y2018F, G2019S, I2020T and phosphorylated at S2032 and T2035 to form the activated LRRK2 kinase (45, 46) and processed in Maestro (Schrodinger). The Protein Preparation Wizard was used to build missing sidechains and model charge states of ionizable residues at neutral pH. Hydrogens and counter ions were added and the models were solvated in a cubic box of TIP4P-EW water (47) and 150 mM KCl with a 10 Å buffer in AMBER tools (48). AMBER16 (48) was used for energy minimization, heating, and equilibration steps, using the CPU code for minimization and heating and GPU code for equilibration. Parameters from the Bryce AMBER parameter database were used for phosphoserine and phosphothreonine (49). Systems were minimized by 1000 steps of hydrogen-only minimization, 2000 steps of solvent minimization, 2000 steps of ligand minimization, 2000 steps of side-chain minimization, and 5000 steps of all-atom minimization. Systems were heated from 0 K to 300 K linearly over 200 ps with 2 fs time-steps and 10.0 kcal·mol·Å position restraints on protein. Temperature was maintained by the Langevin thermostat. Constant pressure equilibration with an 8 Å non-bonded cut-off with particle mesh Ewald was performed with 300 ps of protein and peptide restraints followed by 900 ps of unrestrained equilibration. Gaussian accelerated MD (GaMD) was used on GPU enabled AMBER16 to enhance conformational sampling (50). GaMD applies a Gaussian distributed boost energy to the potential energy surface to accelerate transitions between meta-stable states while allowing accurate reweighting with cumulant expansion. Both dihedral and total potential acceleration were used simultaneously. Potential statistics were collected for 2 ns followed by 2 ns of GaMD during which boost parameters were updated for each simulation. Each GaMD simulation was equilibrated for 10 ns. For each construct 10 independent replicates of 200 ns of GaMD simulation were run in the NVT ensemble, for an aggregate of 2.0 μs of accelerated MD. Potential energy surfaces were determined along each reaction coordinate using a cumulant expansion to the second order.

## Results

To characterize and dissect the functional properties of the catalytic domains of LRRK2, in particular the kinase domain, we used a multi-scale approach that begins with testing real time filament formation in live cells to assessing the consequences of molecular dynamics simulations of PD mutations in the kinase domain. Of primary importance was to characterize the biochemical properties of LRRK2_RCKW_ following deletion of the catalytically inert N-terminal lid. To next confirm that LRRK2_RCKW_ was a well-folded protein we used hydrogen deuterium exchange mass spectrometry (HDXMS) and then to capture the allosteric features of the kinase domain we mapped by HDXMS the conformational changes in LRRK2_RCKW_ that result from adding MLi-2.

### Capturing filament formation in real time

We and others previously showed that treatment with a highly specific LRRK2 inhibitor (MLi-2) induces filament formation of wt LRRK2 and the G2019S mutant when these proteins are transiently expressed in mammalian cells (17, 36). To capture the dynamics of such redistribution we performed time-lapse imaging of YFP-tagged G2019S and wt LRRK2. As shown in **Figure 1** and Supplementary videos, under normal conditions G2019S LRRK2 is mostly diffuse in the cytosol; however, 15-30 min following MLi-2 treatment the protein begins to concentrate first in ‘satellite’ structures diffuse throughout the cells. It then polymerizes to form intricate thicker filaments by 2.0-2.5 h after treatment. Although wt LRRK2 follows a similar redistribution upon treatment with MLi-2, in general it takes longer, approximately 30 min-1 h, before the first structures are observed (Supplementary movie). In both cases this effect is readily reversible: after washout of MLi-2 for 2 h, the proteins gradually diffuse back into the cytosol. To verify that this protein rearrangement was truly dependent on the specific MLi-2 inhibitor, we performed time-lapse imaging using a type 2 inhibitor, rebastinib (**Figure 1B**). Although rebastinib stabilizes a Kinase-WD40 construct of LRRK2, based on a thermal shift assay (**Figure S1**), it did not induce changes in the localization of G2019S proteins even after 8 h treatment, confirming the prediction of Deniston *et al.* (2020) (13).

**Figure 1.**
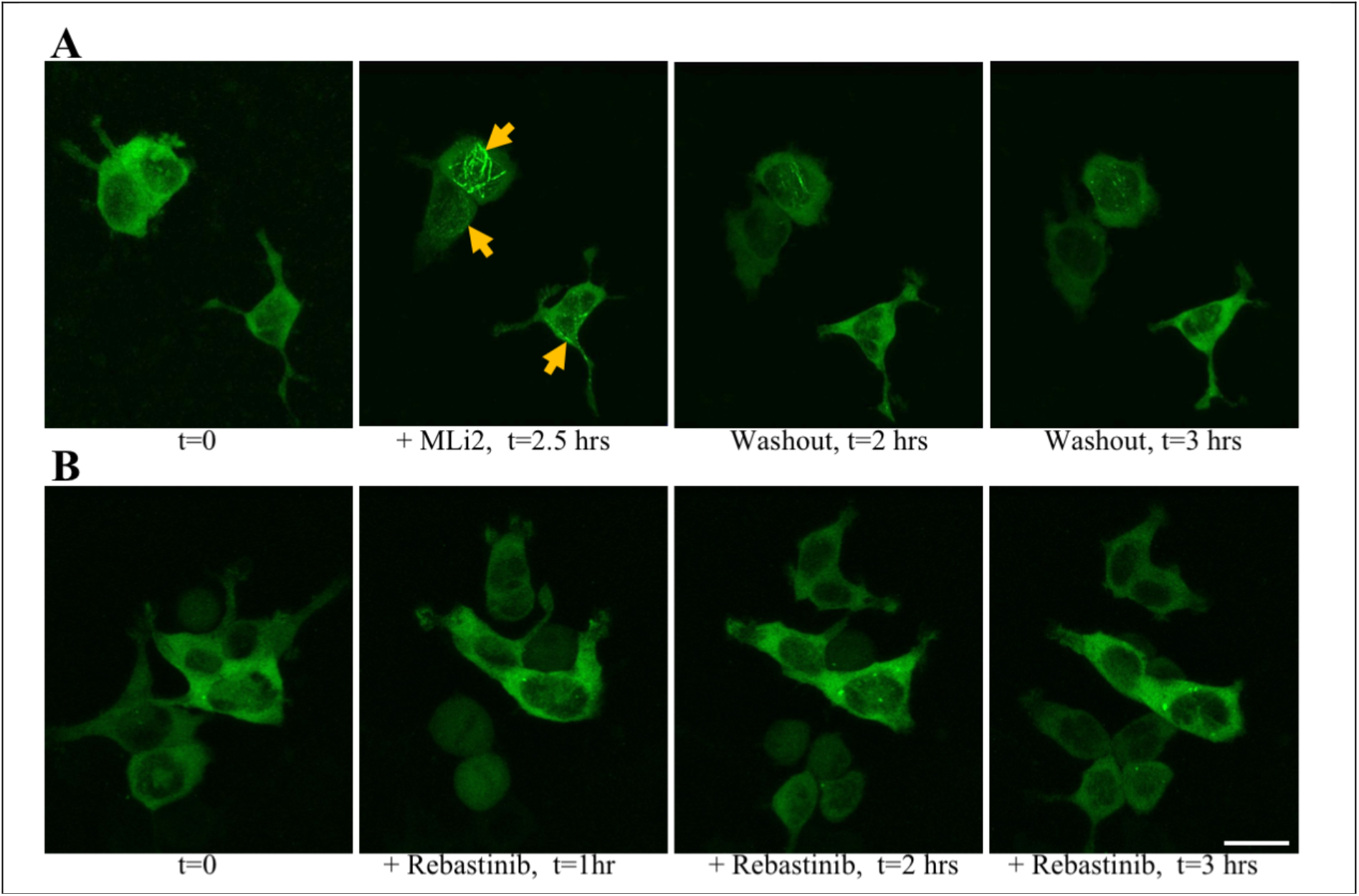
MLi-2, but not Rebastinib, affects the localization of the kinase hyperactive LRRK2-G2019S mutant. **A)** Time-lapse imaging of HEK293T cells transiently expressing YFP-LRRK2-G2019S: confocal images (YFP fluorescence signal, maximum intensity projections) were acquired every 11 min. Representative images show the typical diffuse cellular localization of the proteins (t=0 h) prior to treatment with 100 nM MLi-2; following MLi-2 addition, proteins relocalize to form cytoplasmic filamentous structures (yellow arrows, +MLi-2, t=2.5 h). After washout of the inhibitor, the proteins gradually dissociate from the filaments into the cytosol (washout, t=2-3 h). **B)** Time-lapse imaging of HEK293T cells transiently expressing YFP-LRRK2-G2019S before (t=0 h) and after treatment with 100 nM Rebastinib. No changes in the localization of the proteins are observed. Scale bar, 20 microns.

### LRRK2_RCKW_ variants spontaneously form filaments around microtubules in an MLi-2 independent manner

In our filament formation assay, flag-tagged variants of the LRRK2_RCKW_ construct were overexpressed and cells were analyzed after fixation via antibody staining in a confocal laser-scanning microscope. The majority of the transfected cells, regardless of the mutation, displayed constitutive filament formation (**Figure 2**). Most striking, in contrast to fl LRRK2, is that wt and G2019S are no longer dependent on MLi-2 for docking onto MTs. This supports the hypothesis that the inert N-terminal scaffolding domains are not required for the filaments to form but instead are essential for protecting or shielding the catalytic domains to prevent them from docking onto MTs. In this way they promote the cytosolic distribution of LRRK2 prior to activation, which is likely further facilitated by phospho-dependent interactions with specific 14-3-3 proteins (22, 27). The N-terminal domains are also important for docking to Rab proteins such as Rab29, which are thought to initiate activation of LRRK2 (29). This would be a physiological mechanism where multiple biological functions are embedded in the catalytically inert N-terminal domains, this includes activation and/or localization by heterologous proteins as well as inhibition of the catalytic domains. We hypothesize that most of the PD mutations circumvent or “hijack” this normal process.

**Figure 2.**
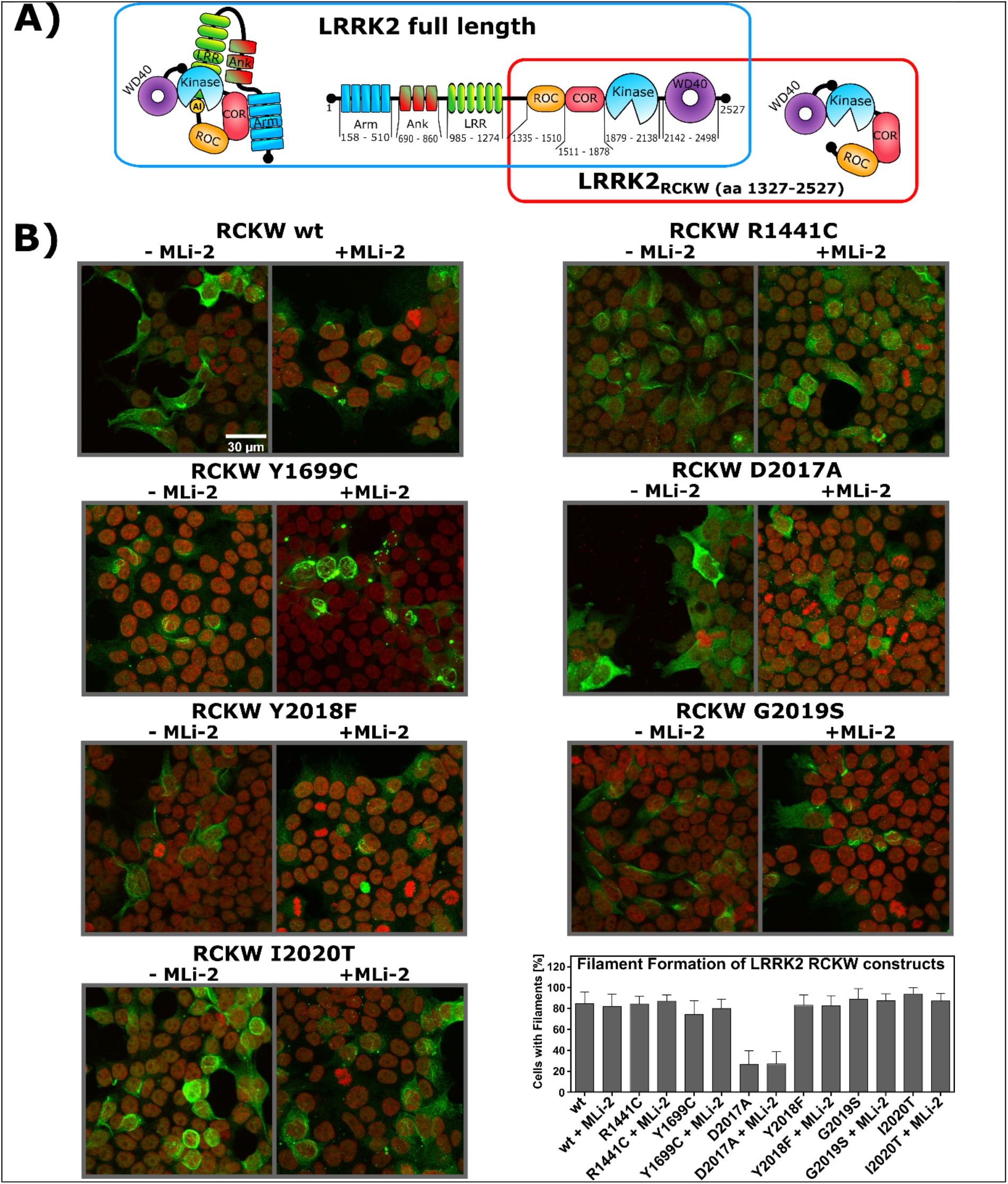
Filament formation of LRRK2_RCKW_ constructs is independent of MLi-2 treatment but reduced in case of LRRK2_RCKW_ D2017A. **A)** Schematic domain organization of LRRK2 full length protein (blue box) and the LRRK2_RCKW_ construct (red box). **B)** Plasmids, encoding LRRK2_RCKW_ variants, were transfected into HEK293T cells for LRRK2_RCKW_ overexpression. Transfected cells were then analyzed for the spatial distribution of LRRK2_RCKW_ by immunostaining. All tested LRRK2_RCKW_ variants displayed a high likelihood (80-90%) to form filaments inside the HEK293T cells except for LRRK2_RCKW_ D2017A (20-30%). Interestingly, in contrast to LRRK2 full length the percentage of cells showing filament formation was independent of MLi-2 treatment or a specific LRRK2_RCKW_ mutation. Scale bar, 30 microns.

Of the mutants tested only LRRK2_RCKW_ D2017A, a kinase dead mutant, showed strongly reduced docking onto MTs, and this is consistent with our earlier findings showing that the full-length D2017A mutant did not dock onto MTs even in the presence of MLi2 (**Figure 2**). We confirmed here that MLi-2 did not have an additive effect on the percentage of cells showing LRRK2_RCKW_ filaments and did not induce binding of the D2017A mutant (**Figure 2**). We conclude that the high affinity binding of MLi-2 to the kinase domain is sufficient to unleash the N-terminal protective lid that normally shields the catalytic domains and promotes localization in the cytosol. We also show that simply removing the N-terminal lid is in most cases sufficient to promote docking onto MTs. The exception is the D2017A mutant, which cannot bind well under any conditions either because it lacks the ability to undergo a subsequent essential auto-phosphorylation step or, most likely, because it is locked into an open conformation similar to what we saw with rebastinib. We next asked whether LRRK2_RCKW_ retained its full kinase catalytic activity even though the regulatory machinery embedded in the N-terminal domains is removed.

### Protein kinase activity is conserved and, in some cases, amplified in the LRRK2_RCKW_ proteins

To assess kinase activity, we used both LRRKtide, a small synthetic peptide, and Rab8a as substrates for the LRRK2_RCKW_ proteins. In addition to wt LRRK2_RCKW_ we measured the kinase activities of two ROC:COR domain mutations (R1441C and Y1699C) and four mutations in the kinase domain, more precisely in the DYGψ motif (D2017A, Y2018F, G2019S, I2020T). R1441 and Y1699 are located in the ROC and COR domains, respectively, and, based on homology models, are predicted to be part of the ROC:COR domain interface (11, 51, 52). Importantly, we found that wt LRRK2_RCKW_ has kinase activity that is comparable to full-length LRRK2 (36) although in the absence of the N-terminal scaffolding domains the activity is no longer dependent on Rab activation. Using LRRKtide as a substrate we found that the pathogenic mutation R1441C slightly increased the kinase activity while Y1699C had only a minor effect on LRRKtide phosphorylation (**Figure 3**). In contrast, when we used a physiological substrate, Rab8a, Y1699C led to an enhanced pT72 phosphorylation *in vitro,* comparable to the phosphorylation by Y2018F, whereas R1441C behaved like wt (**Figure 3**). The fact that kinase activity is dependent on substrate may account for some of the confusion in the literature about the activity of various LRRK2 mutants but suggests that some of the mutations may change substrate specificity. If these residues are indeed at a domain interface, as predicted, they could also introduce a conformational change that would result in the unleashing of the N-terminal scaffolding domains and/or promote dimerization which is associated with membrane localization and activation of LRRK2 (29, 53, 54).

**Figure 3.**
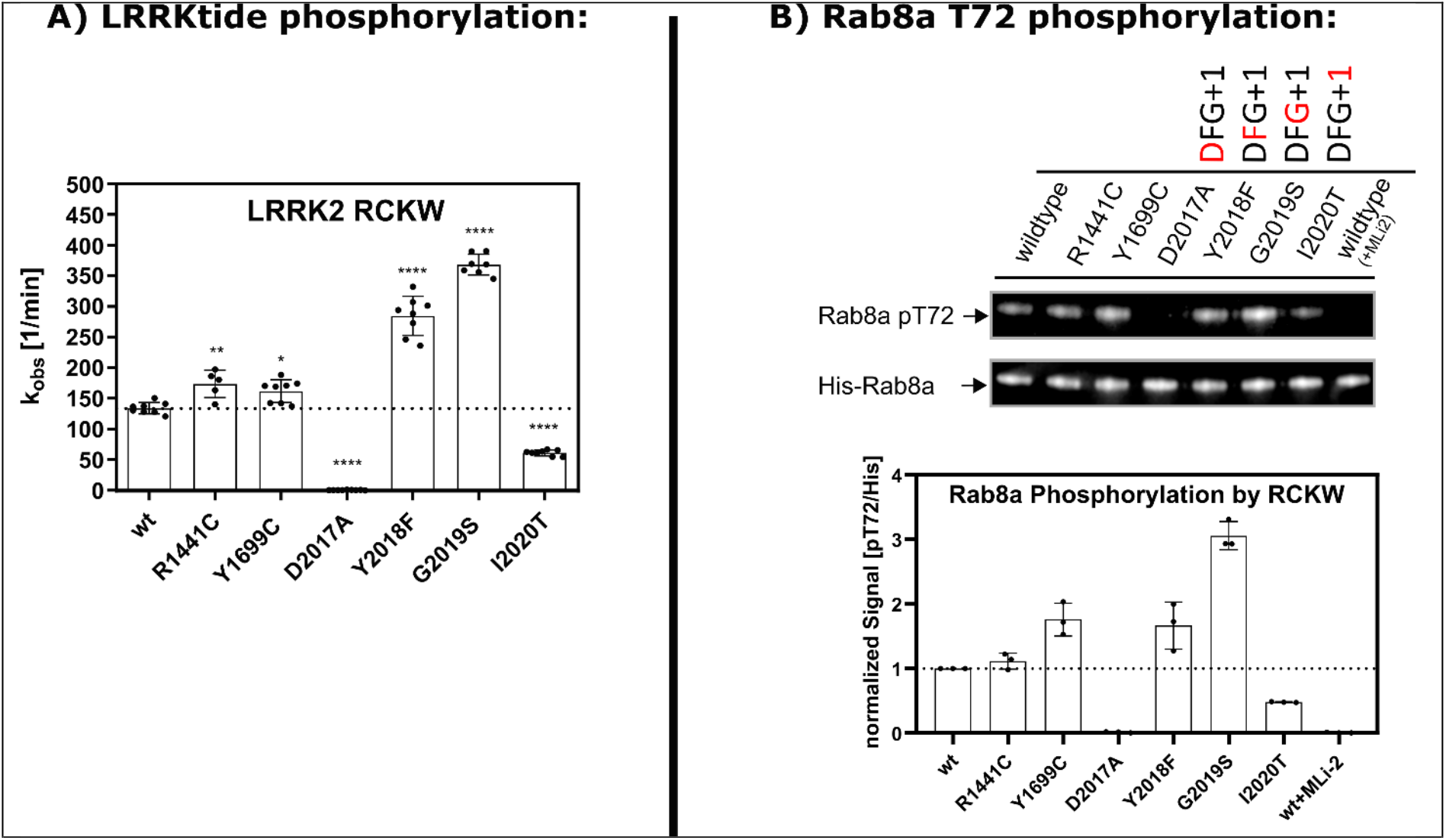
The LRRK2_RCKW_ variants Y2018F and G2019S enhance LRRKtide phosphorylation, while Rab8a phosphorylation is increased by Y1699C and G2019S but not by Y2018F. **(A)** Using the peptide LRRKtide as a substrate for LRRK2_RCKW_ variants revealed that although it is a deletion construct it preserves LRRK2 full length kinase activity. Additionally, the DYGψ mutants tested here also resemble the results of their full-length counterparts. Interestingly, also the pathogenic mutations R1441C and Y1699C which are situated in the RocCOR region of the LRRK2_RCKW_ construct display a mild increase in kinase activity compared to LRRK2_RCKW_ wt. Experiments for each mutant were performed at least in duplicates of duplicates for two independent LRRK2 expressions. Each dot represents the mean of a duplicate, while the dotted line represents the mean of the measured wt activity. To determine significant differences between LRRK2_RCKW_ wt and mutant activity a one-way ANOVA test based on the Dunnett’s multiple comparisons test was performed. Hereby one asterisks (*) indicate a P value between 0.05 and 0.01, two asterisks (**) a P value between 0.01 and 0.001 and four asterisks (****) a P value below 0.0001. **(B)** Besides LRRKtide also a physiological substrate of LRRK2, Rab8a, was tested as a kinase substrate of the RCKW construct. The kinase assays were performed for 7 min at 37 °C and then stopped by adding SDS-sample buffer. Western blotting against the pT72 site of Rab8a and the His-tag of His-Rab8a revealed increased phosphorylation of Rab8a by LRRK2_RCKW_ Y2018F, G2019S and Y1699C. MLi-2 was shown to efficiently block phosphorylation of Rab8a which is also true for the kinase dead mutant D2017A. Quantification was performed for two independent Western Blots using two independent protein preparations. For each quantification, the pT72 signals were referenced to the signal for the His-tag of 6xHis-Rab8a and then normalized to the resulting wt signal. The dotted line therefore represents 100% of the wt signal.

The strongest effects on kinase activity for LRRK2_RCKW_ were observed for mutations embedded within the activation segment of the kinase domain, specifically in the DYGψ motif where ψ is typically conserved as a hydrophobic residue. Most other kinases have a DFGψ motif, and Y2018 was predicted earlier, based on activation when the Tyr is replaced with Phe, to serve as a brake that keeps LRRK2 in an inactive state (36). We measured the effect of mutating each of these residues on kinase activity. The D2017A (DYG) mutant was not able to phosphorylate either LRRKtide or Rab8a (**Figure 3**) which is consistent with other kinases, since this residue is part of the regulatory triad and is crucial for the correct coordination of the Mg^2+^-ions and the γ-phosphate of ATP in the kinase active site cleft (55). Reintroducing the classical DFG motif to LRRK2_RCKW_ (DYG in LRRK2) increases the kinase activity for LRRKtide by a factor of 3-4, whereas Rab8a phosphorylation was only enhanced by a factor of 1.7 (**Figure 3**). LRRKtide phosphorylation by LRRK2_RCKW_ G2019S was comparable to LRRK2_RCKW_ Y2018F. When Rab8a is used as a substrate, G2019S phosphorylation of T72 was two times higher than Y2018F. The other tested pathogenic DYGψ mutation, I2020T, displayed a reduced phosphorylation of LRRKtide as well as Rab8a (**Figure 3**). This is also in accordance with our LRRK2 full-length data of I2020T, albeit full-length I2020T Rab8a phosphorylation was comparable to wt. The results for the I2020T mutation in LRRK2 full length and LRRK2_RCKW_ demonstrate that LRRK2 pathogenicity is not driven solely by increased kinase activity but also by changed substrate preferences such as serine/threonine specificity as well as changes in subcellular localization. We later show that the dynamic properties are also altered by these mutations.

### Mapping the conformational changes induced by MLi-2 using Hydrogen Deuterium Exchange Mass Spectrometry (HDX-MS)

To define the global conformational changes induced in LRRK2_RCKW_ as a consequence of MLi-2 binding we used HDX-MS, which allows us to determine the solvent exposed regions of the protein over a time course of 5 minutes. The exchange data was mapped onto models of the ROC:COR and kinase domains and onto the solved structure of the human WD40 domain. Although this is a large protein (1200 residues), we obtained excellent coverage (>96 %), and the solvent exposed regions are consistent with the predicted folding of all four domains (**Figure S2**). While we focus here primarily on the kinase domain, the graph summarizing the overall solvent accessibility of the entire protein shows not only that the four domains are well-folded but also identifies several regions that are highly solvent exposed. Of particular note is the activation loop of the kinase domain as well as the segment that lies between the COR-B domain and the kinase domain and the segment that joins the GTPase domain to the COR-A domain. The HDXMS data suggests that these regions having high deuterium uptake are highly flexible or unfolded. Conversely, there are also regions on the surface of each domain that are highly protected from solvent, implying that these are domain-domain interfacial surfaces (**Figure S2**). It is important to appreciate that the HDXMS profile is obtained independent of a solved structure and can thus serve as validation of a predicted model. Overall LRRK2_RCKW_ is a well-folded protein that is consistent with a complex topological model with inter-domain interactions.

Under apo conditions the N-lobe of the kinase domain is more shielded from solvent than the C-lobe (**Figure 4A**). The αC-β4 loop, for example, is almost completely shielded from solvent. This is somewhat unusual in that the N-lobe in the absence of nucleotide tends to be rather dynamic for many protein kinases. The ordered and stable structure of the LRRK2 N-lobe is predicted to be due to constraints imposed by the other domains. This is analogous to the way that cyclin binding orders the N-lobe of CDK2 in contrast to the isolated kinase domain (56). Most kinase structures represent just an isolated kinase domain so one cannot appreciate how other domains contribute to stabilization and, in turn, regulation of the N-Lobe. Our HDXMS results also help to explain why it has not been possible so far to express the kinase domain independent of the rest of the protein. In the apo protein the activation loop of the kinase domain in the C-lobe has the highest deuterium uptake suggesting it is highly disordered and exposed to solvent (**Figure 4 and S2**).

**Figure 4.**
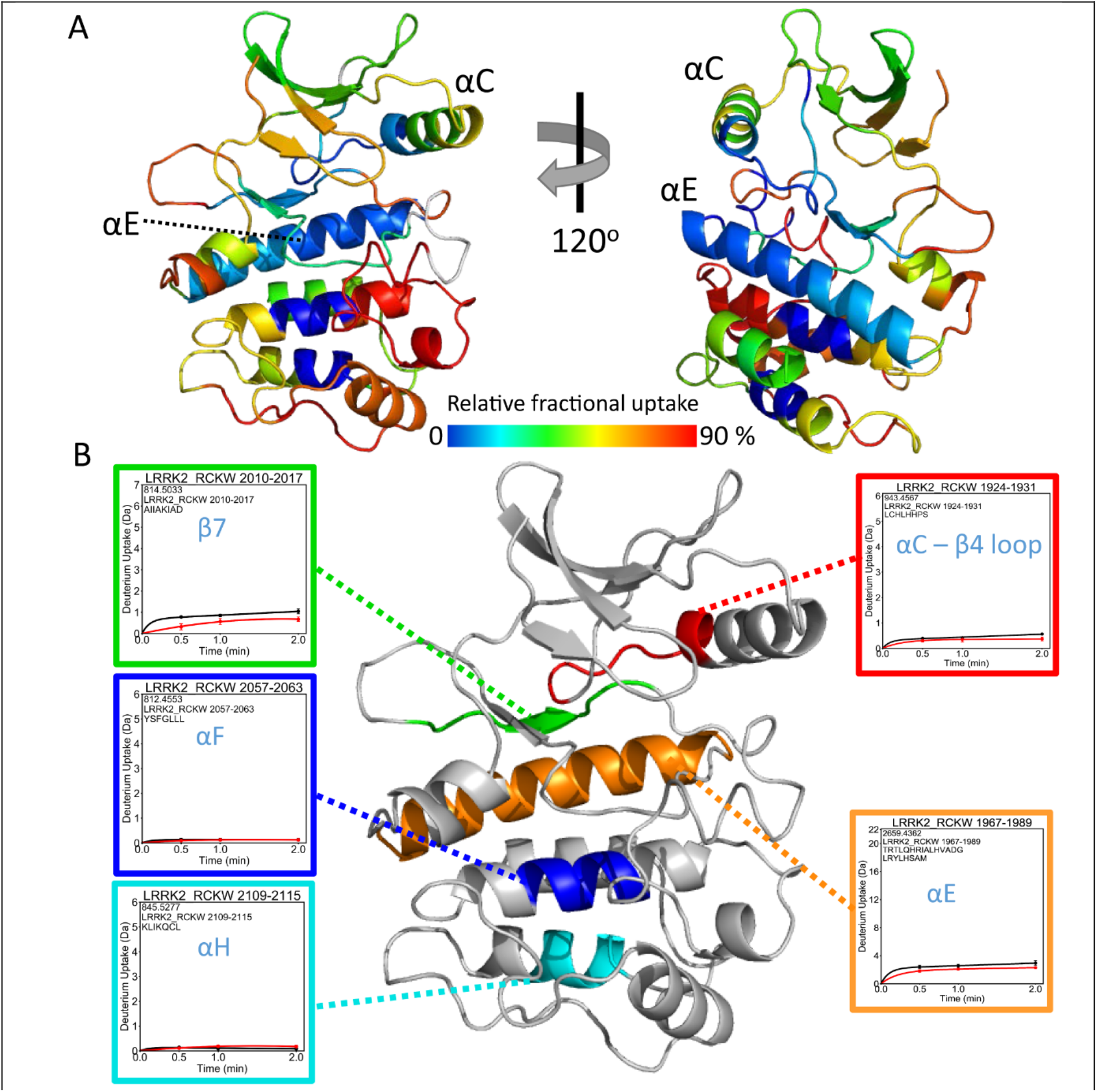
The deuterium uptake of the LRRK2_RCKW_ kinase domain. **(A)** The relative deuterium uptake after 2 min of deuterium exposure of the LRRK2_RCKW_ kinase domain is shown in a color-coded homology model. Grey color indicates no deuterium uptake information. The N-lobe of the kinase mostly shows blue to green colors indicating low deuterium uptake. On the other hand, the αD, activation loop, the end of αF to αH have higher deuterium uptake suggesting a more dynamic, solvent accessible C-lobe. **(B)** Representative peptides that have almost no deuterium uptake are mapped on the kinase domain. Insets show uptake for the apo kinase (black) and the MLi-2 bound state (red).

To gain insight into the allosteric impact of inhibitor binding we next looked at the conformational changes in LRRK2_RCKW_ following treatment with MLi-2. The overall changes, captured in the graph in **Figure 5**, show that there is subtle, albeit important, protection in regions that extend into the GTPase and COR-A:COR-B domains, but by far the largest changes are concentrated in the kinase domain and in the linker that precedes the kinase domain. There are no regions that show enhanced solvent accessibility in the presence of MLi-2. We focus here on the conformational changes that are localized to our kinase domain model. These changes lie not only in the N-lobe and the active site cleft where the inhibitor is directly docked but also in the C-lobe in regions that lie far from the active site cleft.

**Figure 5.**
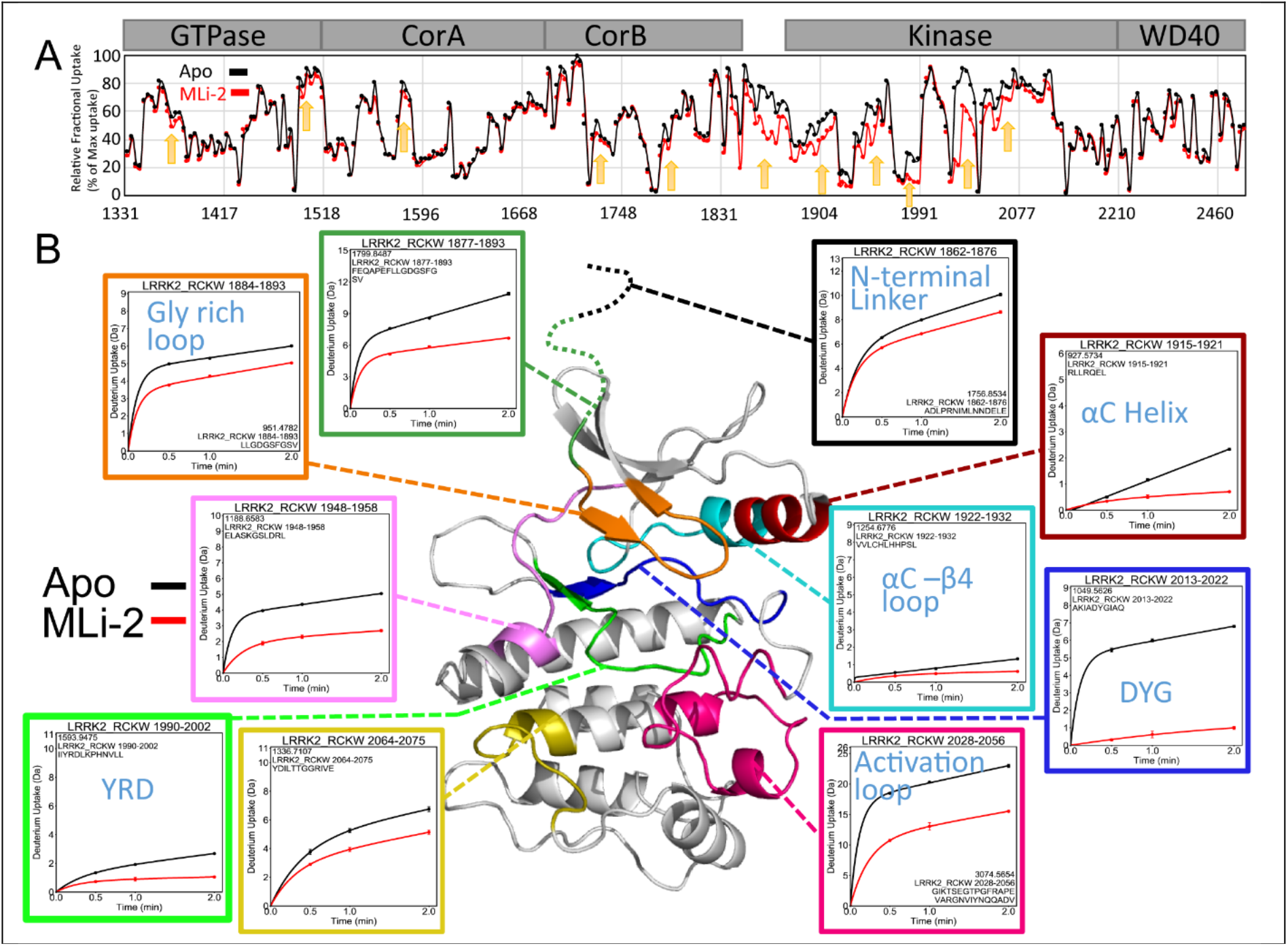
Binding of MLi-2 reduces the deuterium uptake of LRRK2_RCKW_. **(A)** The relative deuterium exchange for each peptide detected from the N to C terminus of LRRK2 in apo-kinase (Black) and MLi-2 bound (red) conditions at 2 min. The arrows indicate the regions of LRRK2 that have less deuterium uptake when bound to MLi-2 **(B)** The deuterium uptake of selected peptides is plotted and mapped on the kinase model. The uptake is reduced in the Gly rich loop, the αC helix, the activation loop, the DYG motif, the YRD motif and the hinge region in the presence of MLi-2.

Looking more carefully at the protected regions, we saw that the binding of MLi-2 reduces the H-D exchange in the ATP-binding site, the activation loop, the αC helix and the hinge region (**Figure 5**). These regions that would be predicted to contact the inhibitor (57, 58) all show significantly reduced deuterium uptake. Peptides, for example, in the hinge region (aa 1948-1958), including the αD helix, experienced a large increase in protection upon MLi-2 binding (50% vs. 20%). The peptide covering the catalytic loop (aa 2013-2022) including the YRD motif also experienced protection (30% to <10%). The Glycine-rich loop (aa 1884-1893) is also highly protected. Most importantly we see that the peptide containing the DYGI motif (aa 2013-2022) is almost completely shielded as a consequence of MLi-2 binding; the deuterium exchange dropped from 70% to less than 10% suggesting that this region, highly solvent accessible in the absence of ligand, becomes almost completely protected by the coordination of the inhibitor (**Figures 5 and 6**). This is quite consistent with the prediction that the kinase domain assumes a compact and closed conformation in the presence of MLi-2. The C-terminus of this peptide contains the beginning of the activation loop, which now appears to be well-folded and shielded from solvent in contrast to the apo structure. Interestingly, the uptake spectra of the two peptides covering the activation loop show an EX1 bimodal distribution, which indicates two different conformations in solution (59). One of these peptides (aa 2028-2056) is shown in Figures 5 and 6. Most other peptides show a single peak indicative of the more typical EX2 exchange kinetics. In addition, MLi-2 treatment also induced slow-exchange in the DYG loop even though it is highly protected. The peptide that covers the N-terminus of the αC helix (aa 1915-1921) shows significant slow exchange even in the absence of MLi-2 that most likely continues beyond 5 min. Although the exchange is quenched in the presence of MLi-2, the slow exchange still persists.

**Figure 6.**
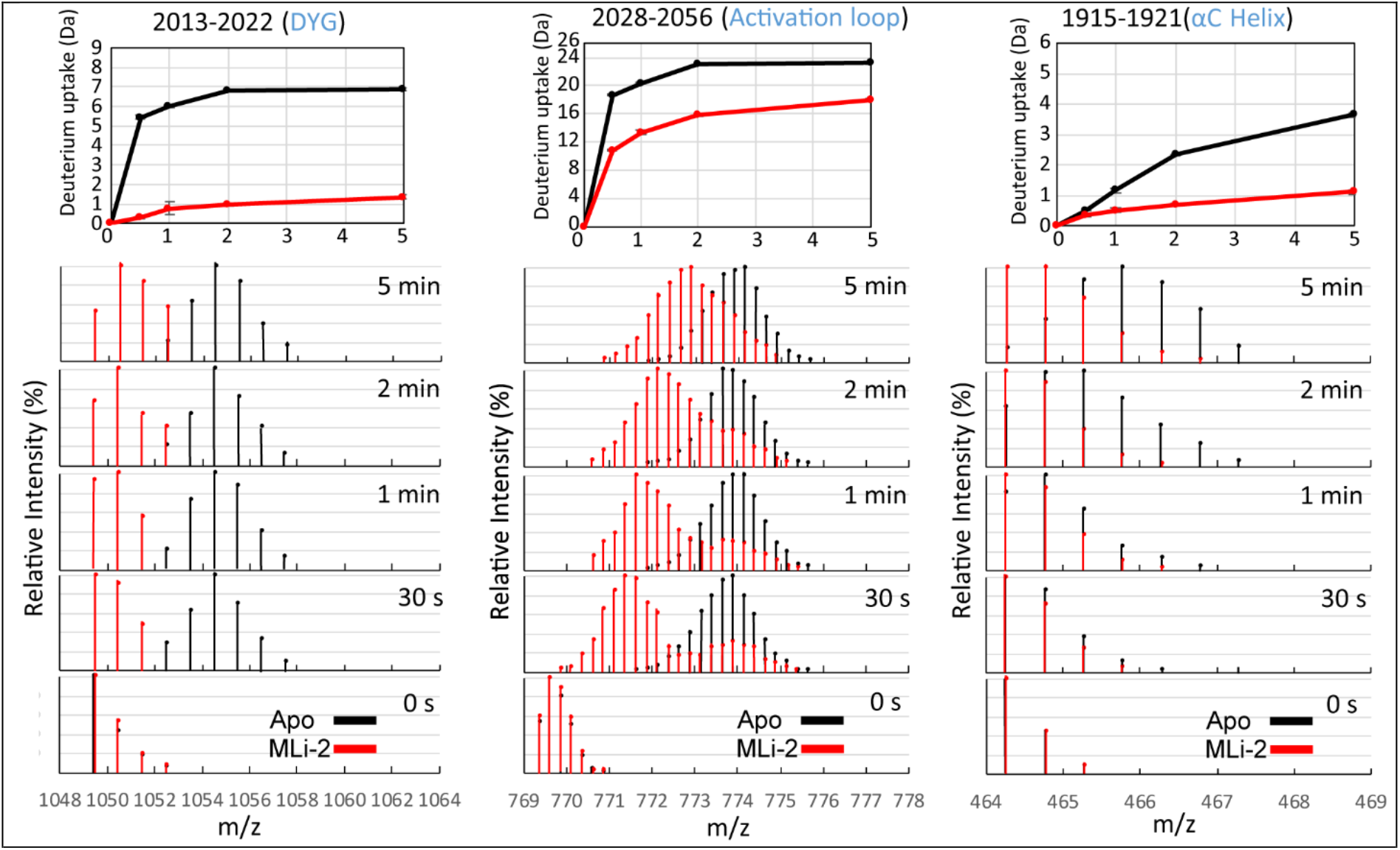
The deuterium uptake and spectral plot of peptides in the DYG, activation loop, and αC helix reveal slow dynamics. In the DYG peptide (2013-2022) the apo state (black) plateaus within 2 min. The MLi-2 bound state (red) continues to slowly exchange at least up to 5 minutes, suggesting that with MLi-2 this region undergoes a slow dynamic process. The apo state of the activation loop peptide (2028-2056) again plateaus within 2 min while the MLi-2 bound state gradually increases after 2 min. From the spectral plot, the uptake of the activation loop peptide in the MLi-2 state exhibits bimodal behavior. One process has slow deuterium uptake (protected); and the other process has fast uptake (solvent exposed)- similar to the single process observed in the apo state. For the αC peptide (1915-1921), the deuterium increases without reaching a plateau over 5 minutes for both states.

The protection of the ATP-binding site and the hinge region by MLi-2 are consistent with other inhibitor bound homolog kinase structures (57, 58) although our data also reflects the dynamic change that the binding of MLi-2 has not only on the kinase domain but also on LRRK2_RCKW_ overall. Essentially any region in LRRK2_RCKW_ that interfaces with the kinase domain will sense binding of nucleotide or an inhibitor. This includes also the LRR domain, not included in our construct, but which is predicted to lie over the kinase domain (13, 51, 60) and would be displaced by the high affinity binding of a kinase inhibitor. HDX-MS shows that changes in conformation and dynamics of the kinase domain are felt through long-distances in LRRK2_RCKW_, as flexible regions throughout the protein exhibit increased protection upon MLi-2 binding (**Figure 5A and S2**).

### GaMD simulations indicate that the LRRK2 kinase domain mutations Y2018F, G2019S and I2020T attenuate flexibility of the activation segment of the kinase core

GaMD simulations were performed on the activated kinase domain of LRRK2 (1865-2135) to investigate changes in the conformational landscape that are caused by the D2017A, Y2018F, G2019S, and I2020T mutations. During all 10 replicate accelerated simulations the wt kinase favors an open and inactive active cleft conformation as measured by the relative position of the N- and C-lobes and the αC-helix (**Figure 7**). The fully closed and active conformation, in which the N- and C-lobes are brought together in concert with an inward positioning of the αC helix to assemble the active site, is infrequently sampled by the wt kinase. In contrast, Y2018F, G2019S, and I2020T are all capable of accessing a closed and active conformation, while D2017A samples a much more open and inactive conformation (**Figure 7a**). The degree of stabilization of the closed conformation roughly correlates with the observed changes in MT association: D2017A<wt<G2019S<Y2018F <I2020T; it does not correlate with activity. The I2020T mutant is trapped in a mostly closed state without extensive open-to-close transitions and αC in-to-out motion compared to wt, Y2018F and G2019S. This loss of breathing dynamics may partially explain the reduced kinase activity of I2020T, where substrate/product kinetics may be impacted. Likewise, Y2018F and G2019S both populate a wide range of open and also closed-active conformations likely contributing to their increased kinase activity. The ability of all of the activating mutants to populate a closed conformation may play a role in their altered MT association compared to wt, where Y2018F and I2020T spontaneously form filaments and G2019S forms filaments faster than wt upon treatment by the type-I inhibitor MLi-2.

**Figure 7.**
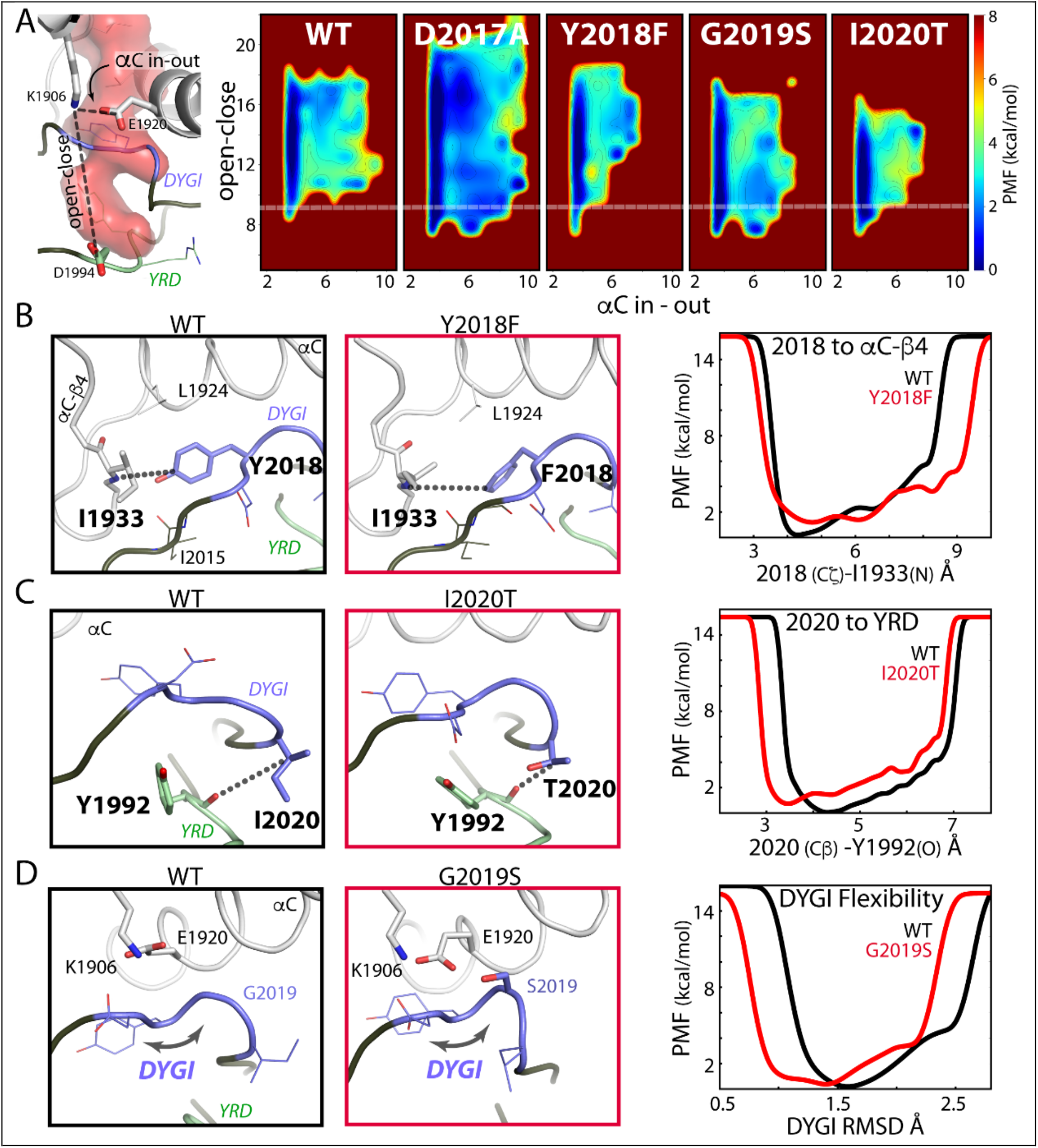
Mutations in the DYGψ loop alter kinase dynamics. **(A)** Kinase conformational free-energy landscape, represented by ‘open-close’: the distance from the top of the active site (K1906/β3 sheet) to the bottom of the active site (D1994/YRD motif), and ‘αC in-out’: the distance between K1906 and the αC helix (E1920). The white dashed line shows the closed-active kinase conformation. The active state is infrequently sampled by the wt kinase, whereas the DYGψ mutants more readily access the closed-active conformation. However, the kinase dead D2017A mutant is destabilized to a more open conformation relative to wt. **(B)** In wt Y2018 (black panel) is locked in an inactive orientation by hydrogen bonds with I2015 and I1933. Y2018F (red panel) packs with L1924 of the αC helix and releases the DYG loop from an inactive state helping to assemble the active site. Y2018F breaks the interaction leading to increased side-chain dynamics, measured by the distance between the 2018 ζ-carbon and the backbone of I2015 (wt:black, mutant:red). **(C)** I2020T makes a hydrogen bond with the backbone of Y1992 in the YRD motif, coupling the DYG and catalytic loops, which results in decreased backbone dynamics. The mutation brings the DYG and YRD motifs together, measured as the distance from the Cβ of 2020 and the backbone of Y1992 (wt:black, mutant:red). **(D)** G2019S bridges the DYG loop to the αC helix and β3 sheet, through E1920 and K1906. This stabilizes the DYG loop, shown by RMSD (wt:black, mutant:red), and promotes the closed kinase conformation.

The hydroxyl moiety of Y2018 in the DYG motif forms persistent hydrogen bonds between the backbone of both I1933 in the αC-β4 loop and I2015 (**Figure 7b**). This interaction stabilizes the tyrosine side chain in an orientation that restricts the αC helix from assembling the active site due to steric clash with L1924. This simulation agrees well with a recent Cryo-EM structure of an inactive conformation of LRRK2_RCKW_, (13). These authors identify the same hydrogen bond between Y2018 and the shell residue I1933. Our simulations provide strong independent evidence that Y2018 in wt LRRK2 is a key stabilizer of the inactive kinase conformation and may also act as a sensor of the αC-β4 loop conformation, a conserved hotspot for kinase allosteric modulation (61). Absence of the OH hydrogen bonds in the Y2018F mutation leads to greater Y2018F side chain dynamics and packing with L1924 that resembles an active kinase configuration (or a properly formed R-spine). An assembled R-spine is the hallmark for an active kinase. Stabilization of the wt Y2018 side chain leads to a ‘frustrated’ DYG backbone free-energy landscape by pulling the motif out of ideal ϕ/ψ space, which is likely important for the kinase’s role as a switch (**Figure S3**). A consequence of freeing the side chain of Y2018F is the convergence of the DYG dihedral angles into a canonically active kinase ϕ/ψ region (62)(**Figure S3,S4**). The I2020T mutation introduces a hydrogen bond between the OH group of the threonine to the backbone of the catalytic YRD motif (Y1992) (**Figure 7c**). The interaction with the YRD motif both stabilizes the catalytic loop and also leads to a closed kinase active site (**Figure 7a,c**). The DYG motif is also stabilized in an active conformation, similarly to Y2018F, as measured by its ensemble DYG dihedral angles (**Figure S4**). The I2020T equilibrium is shifted to the closed conformation and activity may be reduced because the mechanism for opening is impaired. Finally, G2019S introduces a hydrogen bond with the sidechain of E1920 in the αC helix, which in turn forms a highly conserved salt-bridge with K1906 of β3 (**Figure 7d**). The influence of the G2019S mutation on the interaction between αC and β3 and the DYG loop favors the closed and active kinase conformation. The G2019S DYG motif is stabilized in an active conformation as described by its dihedrals (**Figure S4**).

## Discussion

LRRK2 is a highly complex multi-domain protein and equally complex is its regulation. The signaling cascades that control LRRK2 still remain to be elucidated and the molecular mechanisms that control its intrinsic regulation are also not well characterized. However, both aspects are of critical importance to understand the mechanistic consequences of the pathogenic mutations and ultimately find a cure for Parkinson’s Disease. Here we investigated a four-domain construct of LRRK2 consisting of the ROC, Cor, Kinase and WD40 domains (LRRK2_RCKW_). This construct is the shortest functional construct to date that maintains kinase as well as GTPase activity. It is also the smallest construct that can dock onto MTs. In the current work we elucidate different aspects of the intrinsic regulation of LRRK2 using a multilayered approach focusing on the importance of the kinase domain. Our first layer concentrated on the spatial and temporal distribution of full-length LRRK2 in a cellular context as a function of the high affinity kinase inhibitor, MLi-2, which provided us with a real-time assay for filament formation in live cells. The effects of removing the N-terminus on cellular distribution was then explored with our LRRK2_RCKW_ variants. These studies led to the prediction that the N-terminal scaffolding domains shield the catalytic domains in the inactive resting state of LRRK2. The 2nd layer focused on the biochemical characterization of LRRK2_RCKW_ variants by demonstrating substrate-specific kinase activity. Here we showed that the LRRK2_RCKW_ protein retained the catalytic machinery for mediating phosphoryl transfer. In the next layer we used HDX-MS analysis of LRRK2_RCKW_ to provide a portrait of the conformational states of LRRK2_RCKW_ in the presence and absence of MLi2. Mapping the solvent accessible regions in a model of LRRK2_RCKW_ not only provides an allosteric portrait of the kinase domain but also suggests multi-domain crosstalk. Finally, we performed GaMD calculations on the LRRK2 kinase domain to elucidate at a molecular level the differences in breathing dynamics between LRRK2 wt and the pathogenic kinase domain mutations Y2018F, G2019S and I2020T. With this approach we were able to clearly demonstrate that the kinase activity and the spatial distribution of LRRK2 is not only regulated by single domains but by a complex interplay of all the embedded protein domains. The highly dynamic kinase domain, nevertheless, seems to play a crucial role in coordinating the overall domain crosstalk and serves as a central regulatory hub for the intrinsic regulation of LRRK2.

### Filament formation is dependent on unleashing the catalytic domains and on the conformation of the kinase domain

Although multiple functions are associated with the many domains of LRRK2, these domains can be structurally and functionally divided into the catalytically inert NTDs and the catalytic CTDs, and distinct functions are embedded in each. Further complexity is introduced by heterologous proteins such as Rab GTPases and 14-3-3 proteins, which also contribute to the activation and subcellular localization of LRRK2. LRRK2 also exists in multiple oligomeric states where the most active state is thought to be a dimer in contrast to the less active monomer (53, 54). The stability of the monomers and dimers can be further facilitated by heterologous proteins, in particular, the 14-3-3 proteins. Physiologically LRRK2 is thought to be activated by Rab GTPases such as Rab29, which dock onto the NTDs and target LRRK2 to organelles such as the trans Golgi network (29). Other Rabs may also activate LRRK2 but target it to different organelles (28, 63) while auto-phosphorylation on residues such as S1292 likely are subsequent steps in the activation process (13, 64). Many of the PD mutations “hijack” these finely tuned regulatory mechanisms. Constitutive localization to MTs is a phenotype displayed by three of the four common LRRK2 PD mutants (R1441C, Y1699C and I2020T) while the hyperactive G2019S mutant, like wt LRRK2, remains cytosolic. Using cryo Electron Tomography (cryoET) Watanabe and co-workers (60) were able to capture the precise way in which a LRRK2 mutant (I2020T) can polymerize and dock onto MTs. They show specifically how the I2020T LRRK2 mutant forms periodically repeating dimers, which then polymerize in a helical array onto MTs. The cellular phenotypes associated with G2019S, I2020T, and R1441C/Y1699C and other PD associated mutations include perturbation of MT-related processes such as vesicular trafficking, autophagy, cilia formation, and nuclear/mitochondria morphology, so it is very likely that LRRK2 dysfunction physiologically interferes globally with dynamic cross talk with MTs (20, 32, 65–72).

With live cell imaging in the absence and presence of a kinase inhibitor, MLi-2, we were able to capture at low resolution in real time the re-localization of cytoplasmic wt and G2019S LRRK2 to decorated MTs. MLi-2 binding to wt and G2019S thus introduces artificially the pathogenic phenotype constitutively observed for I2020T, R1441C or Y1699C. In contrast to MLi-2 and LRRK2-IN-1, which are type I kinase inhibitors, we show that in the presence of a type II kinase inhibitor, rebastinib, G2019S LRRK2 remained cytosolic confirming the predictions of Deniston and coworkers (2020) (13), that docking to MTs is extremely sensitive to the conformational state of the kinase domain. Although the MLi-2 complex is catalytically inactive, MLi-2, which is a competitive inhibitor of ATP, nevertheless locks the kinase into an active-like conformation (58). In contrast, rebastinib is likely to stabilize a DYG/DFG-out/open and inactive conformation of the kinase domain. In the absence of the NTDs wt and G2019S LRRK2_RCKW_ spontaneously form filaments independent of MLi-2. Our HDX-MS data confirms that deletion of the NTDs in LRRK2_RCKW_ does not unfold the remaining CTDs as solvent exchange shows that the protein is properly folded with the domains well packed against each other. Collectively our results support a model where the catalytically inert NTDs function as a “lid” that shields the active sites of the CTDs. The lid can be unleashed physiologically by activating Rab GTPases or by mutations that make LRRK2 a risk factor for PD. Two of the PD mutants, R1441C/G/H and Y1699C, are localized in the ROC and COR domains, respectively, and presumably disrupt a domain-domain interface which is sufficient to unleash the NTDs. While targeting, activation, and inhibition of the CTDs are important functions that are embedded in the NTDs, our biochemical studies here show that all of the kinase activity of the full length LRRK2 is embedded in LRRK2_RCKW_. The other two common PD mutations (G2019S and I2020T), although functionally distinct, are in the kinase domain. Our live cell imaging together with our biochemical and simulation results, discussed below, and a recently published cryoET structure (60), all support the hypothesis that unleashing the NTD lid as well as an active conformation of the kinase domain, not necessarily kinase activity, are essential requirements for dimerization and MT binding.

### The switch mechanism for activation of LRRK2 is embedded in the DYGψ motif of the kinase domain

With HDXMS we confirm that a shift in conformation is induced by the binding of MLi-2 to LRRK2_RCKW_ and, significantly, we find that stabilizing the active kinase conformation by MLi-2 drove changes in conformation and domain organization throughout LRRK2_RCKW._ The global decrease in backbone deuterium exchange measured across all four domains particularly in the linker between the CORB and kinase domains and in flexible regions throughout LRRK2_RCKW_ (**Figure 5A**) is suggestive of changes in domain:domain packing as well as changes in global conformational dynamics. We propose that changes in the CTDs organization and dynamics, driven by the stabilization of the active kinase conformation, are likely coupled with association and dissociation of the NTDs. This explains why mutations such as R1441C and Y1699C, that lie far from the kinase active site but at a domain interface are capable of unleashing the NTDs. Using the MLi-2 bound LRRK2_RCKW_ as a proxy for the “frozen” closed and active-like conformation of the kinase domain, we explored each of the activating DYGψ mutants using MD simulations and asked how each mutation perturbs the conformational ensemble of the kinase domain.

Our first hint that the DYGψ motif impacts the kinase conformation with consequences on LRRK2 global conformation and regulation came from our previous work with the DYGψ mutations, Y2018F and I2020T, where we were able to correlate changes in the internal organization of the kinase domain, specifically assembly of the Regulatory Spine, with *in-situ* MT association (36). Our data here, shows that deletion of the NTDs induces a similar MT association phenotype independent of mutation. This further confirms that filament formation of Y2018F and I2020T in full length LRRK2 is based on unleashing of the NTDs caused by perturbation of the conformational dynamics of the kinase domain. Although full-length G2019S, the most prevalent PD mutation, does not form significant filaments spontaneously, we show that its kinetics of MT association are increased relative to wt following treatment with MLi-2 (**Supplementary Movie**), suggesting that this mutation may share a common de-regulating mechanism through changes in the kinase domain conformation. Indeed, our GaMD calculations show that stabilization of the DYGψ dynamics by all three mutations Y2018F, G2019S, and I2020T promotes the active kinase conformation. Whereas previous metadynamic studies showed a stabilization of the “DYG-in” over the “DYG-out” conformation (73, 74), we observe in our unbiased MD for Y2018F, G2019S, and I2020T a subtle coalescence of DYG dihedral angles into an active-DYG loop conformation (**Figure S4**). However, each of the mutants favor the active conformation by different mechanisms. Y2018F releases the sidechain from an inactive non-preferred rotamer with frustrated DYGψ backbone dynamics leading to closure by packing with L1924 of the αC helix. In contrast, G2019S connects and stabilizes the DYG motif through hydrogen bonding to the activating αC-β3 salt-bridge (interaction of K1906 and E1920), while I2020T at the ψ position stabilizes itself with the YRD (Y1992, R1993 and D1994) catalytic motif through hydrogen bonding. Significantly, the wt kinase in these simulations still fluctuates between both inactive and active states while favoring the inactive conformation. This means input from external factors, such as the NTDs, are required to fully modulate the conformational equilibrium to inhibit kinase activity. Activating DYGψ mutations subvert the built-in regulation by distorting the inherent conformational equilibrium to a degree that breaks these layers of control. On the other hand, the kinase dead D2017A which has significantly reduced localization to MTs even in the absence of the NTDs cannot be stabilized in an active-like kinase conformation because its active-site cleft is dynamically destabilized. Together with our findings for rebastinib and MLi-2 this further emphasizes that filament formation is not solely dependent on the inhibitory lid-function of the NTD but also on the kinase domain integrity/conformational state.

Our findings highlight that the activation of LRRK2, while simplistically represented by static conformations (i.e. simple active and inactive conformations), is more accurately defined in terms of a shift in conformational ensembles and associated dynamics. This is illustrated not just in the MD simulations by changes in bulk conformations due to DYGψ mutants but also by changes in time-scales of dynamics highlighted by live cell imaging and by the induction of bimodal HDX kinetics after binding MLi-2. HDX shows that the activation loop peptide has two distinct conformational populations. One represents a minor population of a highly solvent exposed species similar to what was seen in the apo state and an additional highly protected species, which gradually becomes more solvent exposed (**Figure 6**). In MD simulations the DYGψ activating mutants mirror this behavior and shift the equilibrium towards an ordered and less solvent accessible activation loop (**Figure S5**). MLi-2 binding appears to lead to an even greater shift in this equilibrium and a large decrease in the kinetics of the exchange between states, effectively trapping a major population of a closed and active-like state of the kinase domain. Extending this concept of regulation by tuning of conformational dynamics to the cellular level, our work implies that changes in the balance of LRRK2 conformational equilibrium will lead to proportional changes in its cellular distribution, i.e. the population of LRRK2 in the cytosol vs. associated with MT should mirror the conformational distribution of expanded and closed LRRK2.

### LRRK2 and BRaf share the same Kinase Activation Mechanism

By comparing the resulting finely tuned multi-layered regulation mechanism of LRRK2 with other related homologs of the kinase tree we recognized that our model for LRRK2 regulation closely resembles the activation process of another multi-domain kinase: BRaf (75–77). LRRK2, like BRaf, is activated by the interaction of its N-terminal non-catalytic domains with a small GTPase: Rab vs. Ras. In both cases autoinhibitory sites/domains (AI) in the NTDs become displaced when the activated GTPase binds, and this unleashes the kinase domain (**Figure 8)**. The kinase domains, no longer locked in their inactive conformations, are free to toggle between their inactive and active states. Only in the active conformation are the catalytic domains able to dimerize. In BRaf this pushes the kinase domain into the active conformation and induces cis-autophosphorylation of the activation loop. Dimerization of the catalytic domains of LRRK2, as seen in the cryoET structure (60), also requires an active conformation of the kinase domain, although in the case of LRRK2 the dimer interface most likely does not directly involve the kinase domain. In both cases this last step stabilizes the kinase domain in a conformation where the R-spines are assembled, rendering them ready to bind and phosphorylate their substrates. When this process is disrupted by a mutation such as Y2018F or I2020T in the kinase domain of LRRK2, kinase activation becomes independent of Rab binding, as these mutations shift the equilibrium to a more active kinase conformation which also promotes displacement of the NTDs (**Figure 9A**). This recapitulates a theme observed in BRaf, wherein the most activating cancer-driving mutation, V600F, renders the kinase active without the need for heterologous activation by Ras (78).

**Figure 8.**
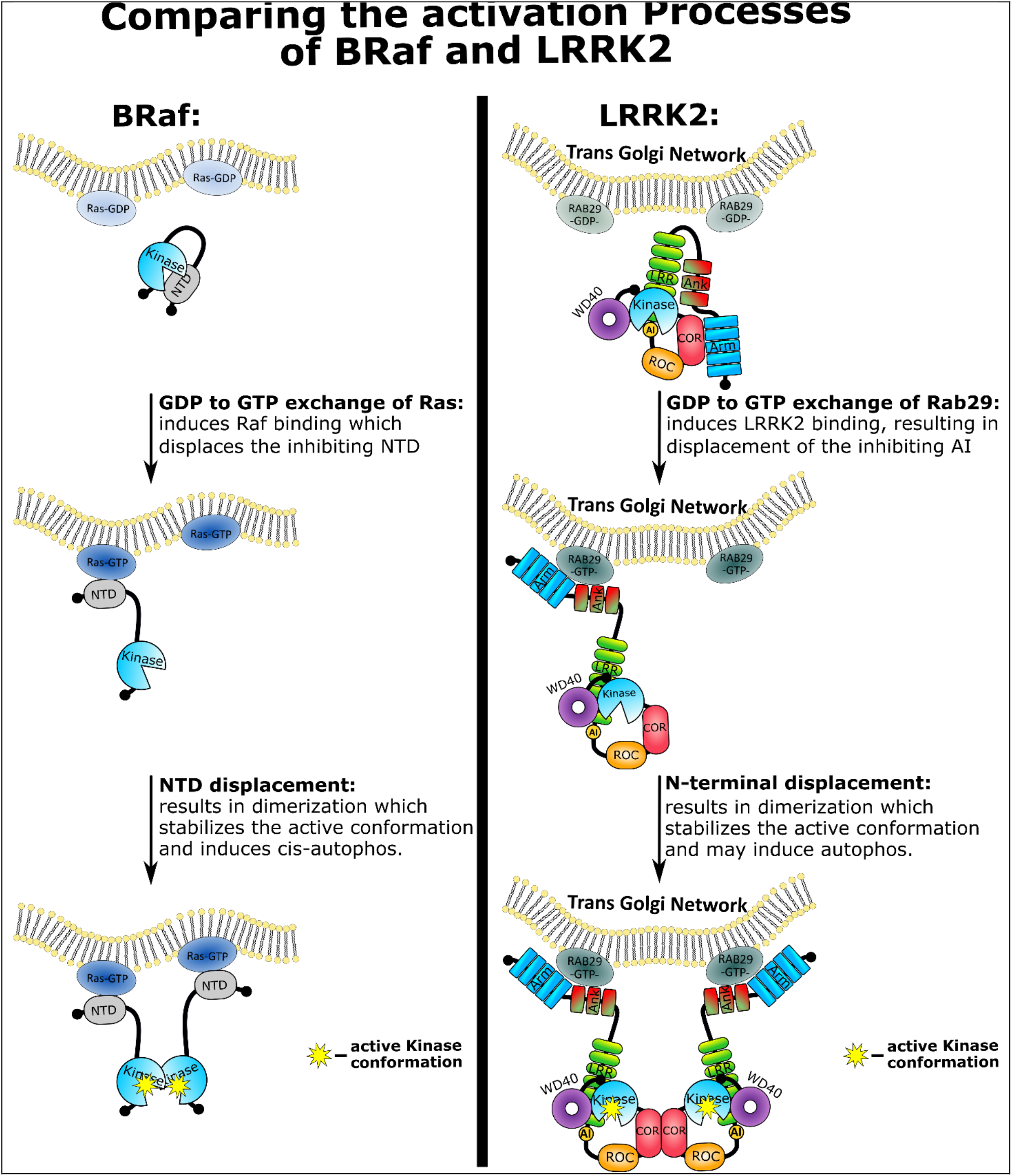
Comparison of the BRaf and LRRK2 activation models. Under healthy conditions both BRaf and LRRK2 are in inactive autoinhibited states in the cytosol. The kinase activation then follows recruitment by either GTP bound Ras or Rab29, which are both small GTPases and closely related to each other. The recruitment induces a conformational change which displaces the N-terminal domain (NTD) of BRaf or the autoinhibition site (AI) in the NTDs of LRRK2. This in turn allows both kinases, BRaf and LRRK2, to dimerize, resulting in stabilization of the active kinase domain conformations. Thereby, the kinases become activated as *cis*-autophosphorylation of the activation loop is induced for BRaf and is also likely to happen for LRRK2. Finally, this unleashes both kinases and result in maximal activation, allowing for efficient phosphorylation of their substrates.

**Figure 9.**
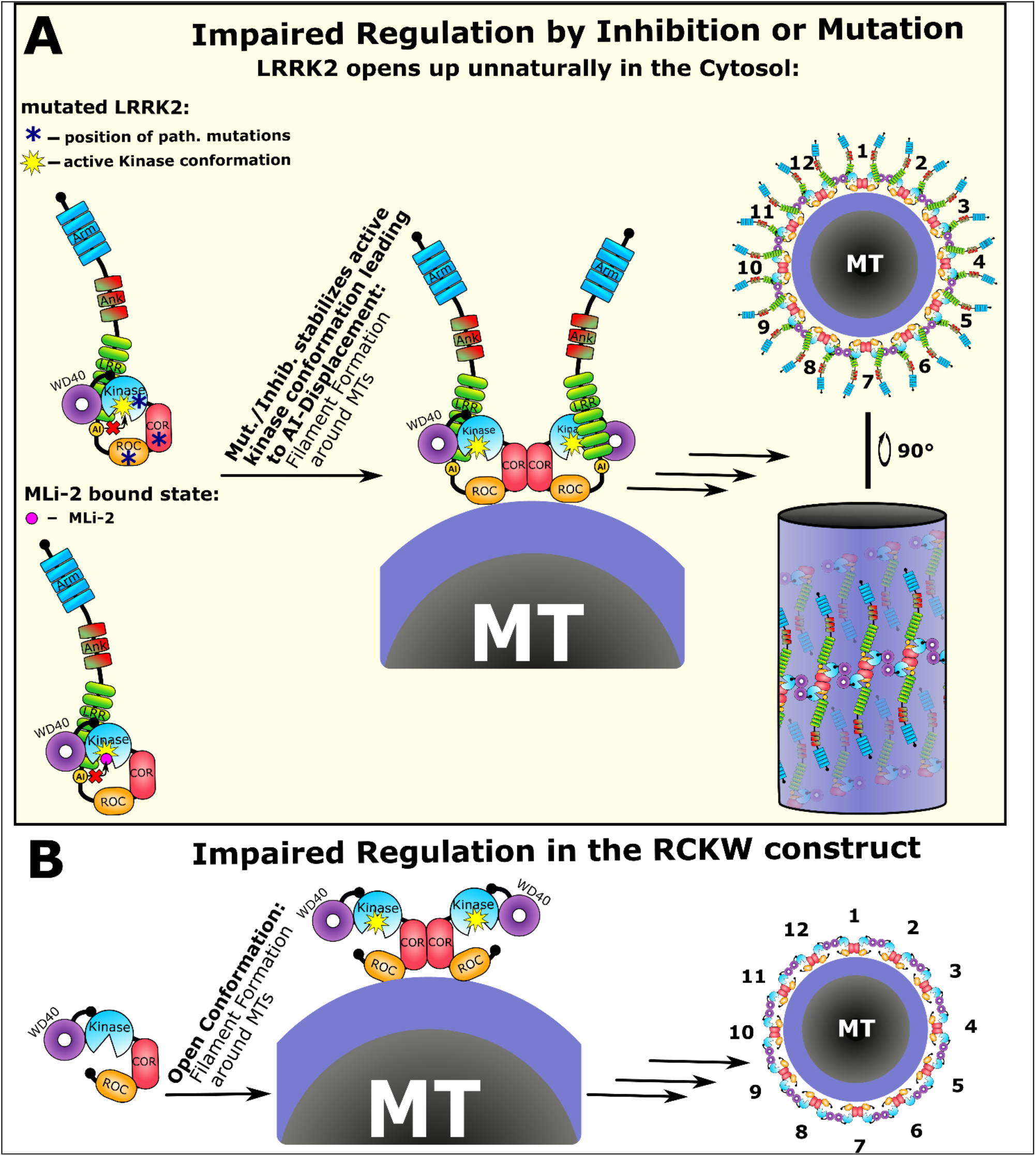
Impaired LRRK2 activation results in microtubule association. **A)** In the mutated or MLi-2 inhibited state the autoinhibitory site (AI) in the NTDs becomes already displaced in the cytosol and the kinase domain is in a constitutively active conformation. This short circuits the activation process which depends on Rab29 association. The active kinase conformation is sufficient to induce dimerization and thereby multimerization of LRRK2 around MT. Using MTs as scaffolding structures for LRRK2 multimers, which are ordered in a periodic fashion as shown by Watanabe and coworkers (2020), result in the filament formation phenotype (60). Each LRRK2 monomer provides two interaction surfaces namely the COR and the WD40 domain which allows each LRRK2 monomer to interact with two adjacent monomers. An additional finding was that the N-terminus forms bridging interactions with the upper and the lower turn of the LRRK2 filament. Furthermore, the number of LRRK2 dimers needed for one turn correlates well with the number of MT protofilaments. Therefore, we believe that this is a pathogenic but specific interaction with MTs resulting from impaired regulation of LRRK2 by stabilizing an active conformation of the kinase domain of LRRK2 either by mutations or by binding of MLi-2. **B)** The importance of the NTDs for stabilizing LRRK2 in an inactive cytosolically distributed state was tested by deleting the N-terminus. The resulting LRRK2_RCKW_ deletion construct spontaneously forms filaments around MTs and is no longer able to become recruited by Rab29 as it lacks the interaction sites in the Arm and/or Ank domain. We believe that the missing AI domain, which we think is positioned in the NTDs, forces the kinase domain of the RCKW construct into an active conformation resulting in multimerization and docking to MTs.

Other mutations that contribute to stabilization of the assembled R-Spine also lead to constitutive activation that is independent of Ras (78, 79). Kinase inhibition was also shown to facilitate downstream signaling as the inhibition of BRaf stabilized the active kinase conformation which is sufficient to promote *cis*-autophosphorylation through heterodimerization (80, 81). This closely resembles the situation we observed for LRRK2 where MLi-2 stabilizes the active kinase conformation and thereby induces filament formation (**Figure 9A**). Another feature common to LRRK2 and BRaf is that the deletion of the N-terminus of either protein generates a constitutively active kinase (**Figure 9B**) (82–84). Finally, the kinase dead mutations in both proteins are unable to form productive dimers (85, 86). Since many aspects of the kinase domain regulation of LRRK2 and BRaf appear to be quite similar, it can be concluded that activation mechanisms, in general, where the kinase domain switching between active and inactive conformations serves as a central hub are likely to be conserved throughout much of the kinome. In addition, from the recent structures of full length BRaf in complex with its substrate MEK and 14-3-3 proteins we can see how these auxiliary proteins can stabilize either an inactive or an active conformation (87). This will certainly also be true for LRRK2. Clearly, we can learn much about LRRK2 activation by looking at close homologs that have been studied for a longer time, and hopefully this will facilitate the discovery of new therapeutic strategies for attacking LRRK2 as a driver of PD.

## Conclusion

Analysis of multidomain kinases suggest that the conformation of the kinase domain might, as a general principle, regulate much more than just the activity of the kinase. Our results with LRRK2 demonstrate that the active kinase conformation not only switches the kinase into an “on” state but also unleashes inhibitory domains, can promote dimerization, and can facilitate translocation to anchored substrates. In the case of LRRK2 this is a complex and tightly regulated process where more domains and other proteins than just the kinase domain are involved; however, the paradigm is similar for BRAF and probably for most kinases such as Src and PKC that are also embedded in multi-domain proteins. In each case, the conformation of the kinase domain seems to play a crucial role in these intrinsic regulatory processes. We also further confirm here how a switch mechanism for activation, embedded in the DYGψ motif of the kinase core, allows the kinase to toggle between inactive and active conformations which is then communicated to all parts of the protein. We also demonstrate here that the non-catalytic NTDs play an important regulatory role by shielding the catalytic CTDs in the absence of physiological activators. Interacting proteins like 14-3-3 or Rab proteins are likely to further fine-tune this regulation either positively or negatively by stabilizing certain conformations of LRRK2. This precisely controlled signaling process can also be hijacked by a variety of disease driving mutations such as those that lead to PD.

## Acknowledgments

We like to acknowledge the excellent technical assistance of Michaela Hansch, Irmtraud Hammerl-Witzel and the assistance of Alexandr Kornev in preparation of models.

## Supporting information

**Figure S1.**
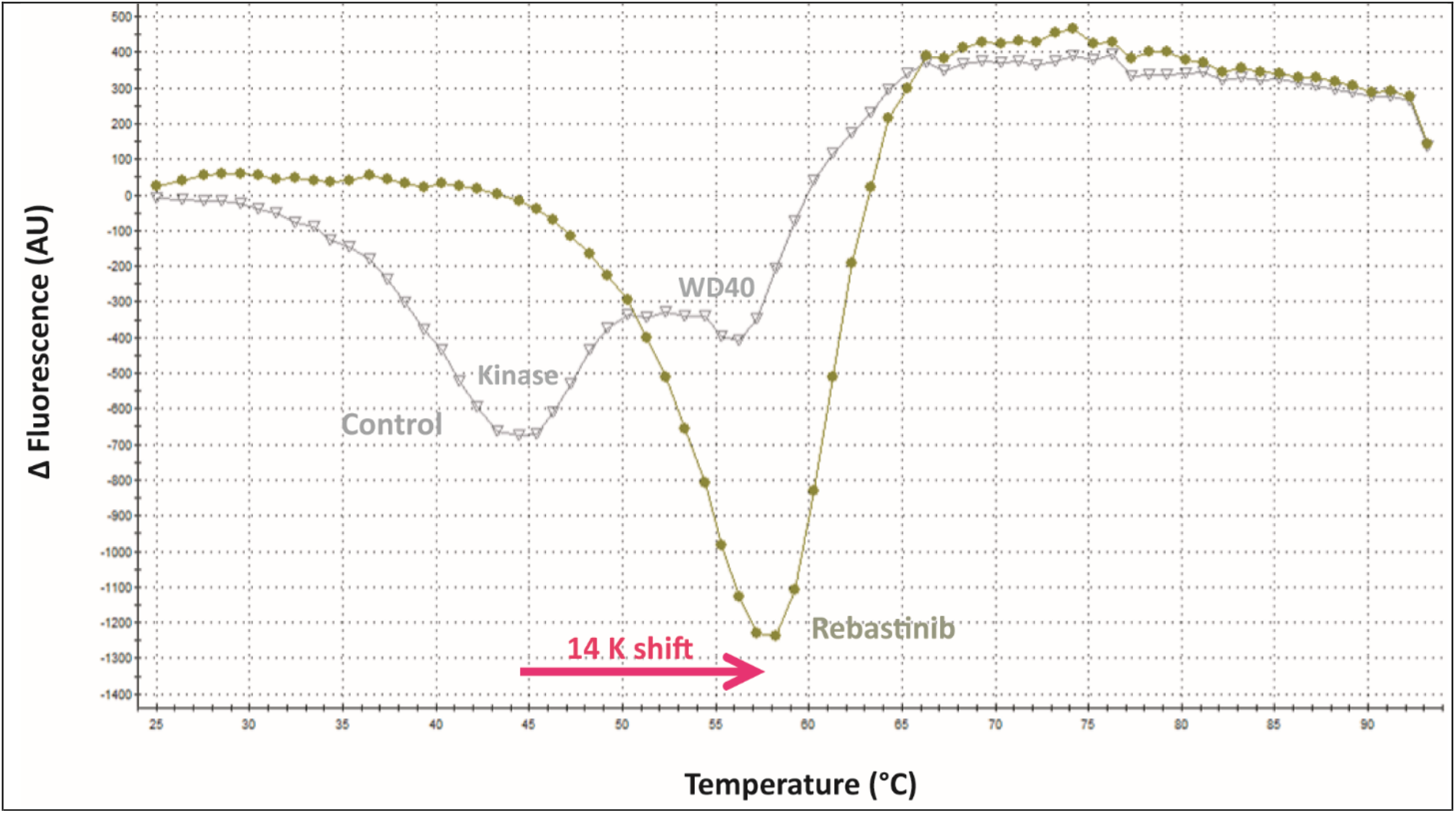
The type-2 kinase inhibitor rebastinib binds to the LRRK2 kinase domain. Shown are the 1st derivatives of the LRRK2 melting curves with or without rebastinib. The LRRK2 protein comprised two domains (kinase and WD40). Accordingly, there were two minima in the control curve. The addition of rebastinib stabilized the LRRK2 kinase domain and shifted its melting temperature by 14 K, indicating a binding constant <1 uM.

**Figure S2.**
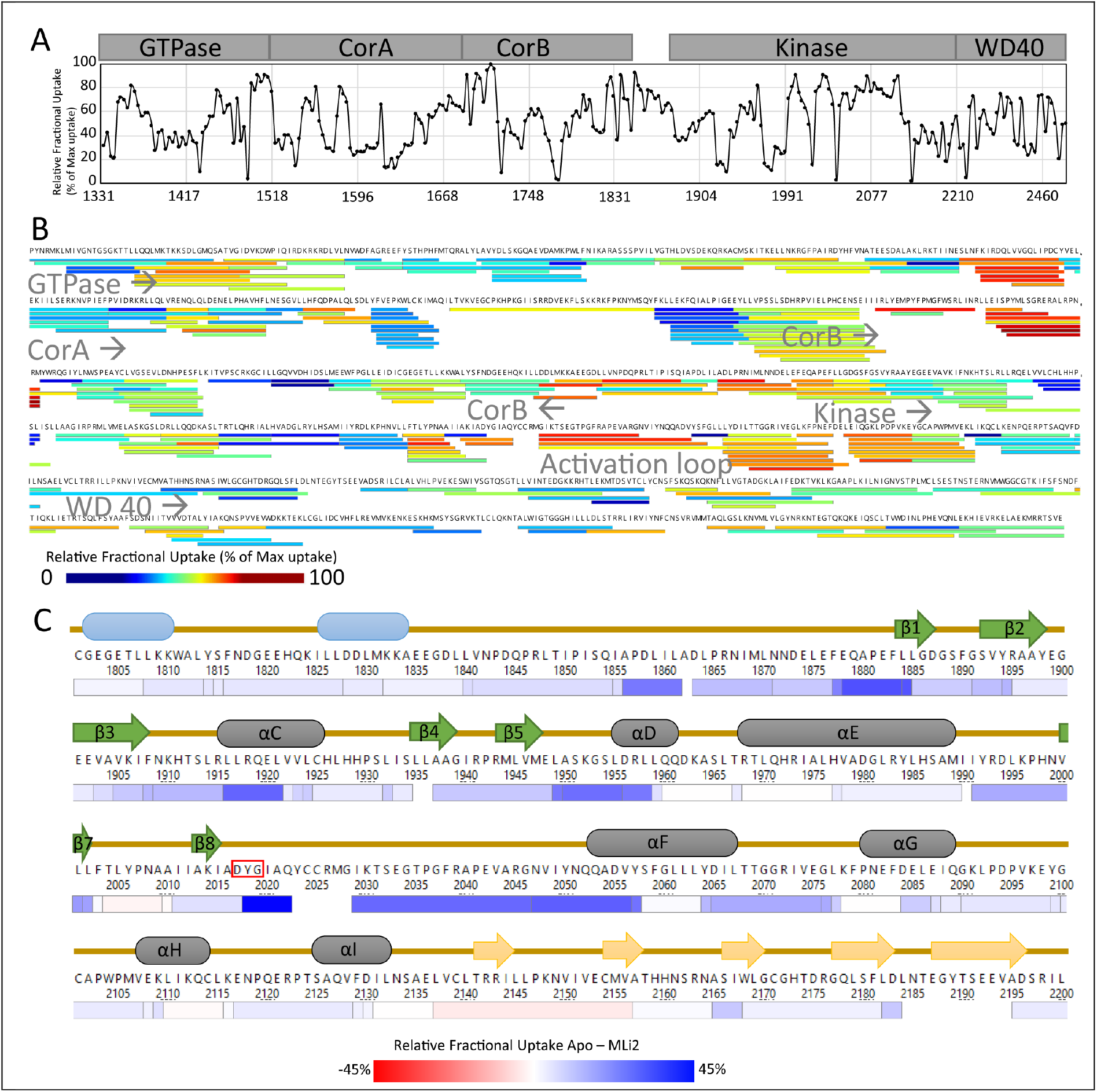
The identified HDX-MS peptides of LRRK2_RCKW_. **(A)** The relative deuterium exchange for each peptide detected from the N to C terminus of LRRK2 at 2 min, apo condition. **(B)** Each line in the coverage map represents an identified peptide (exchanged 2 min) and the color indicates the relative deuterium uptake. Location of each domain is indicated. The map identifies folded regions as well as solvent exposed regions such as the activation loop in the kinase domain. The coverage is 96.3% and the redundancy is 3.72, which is successful for a protein this large. **(C)** The heat shows the relative fractional uptake by color at 2 min. It is colored based on the difference of relative deuterium uptake between apo and MLi-2 states.

**Figure S3.**
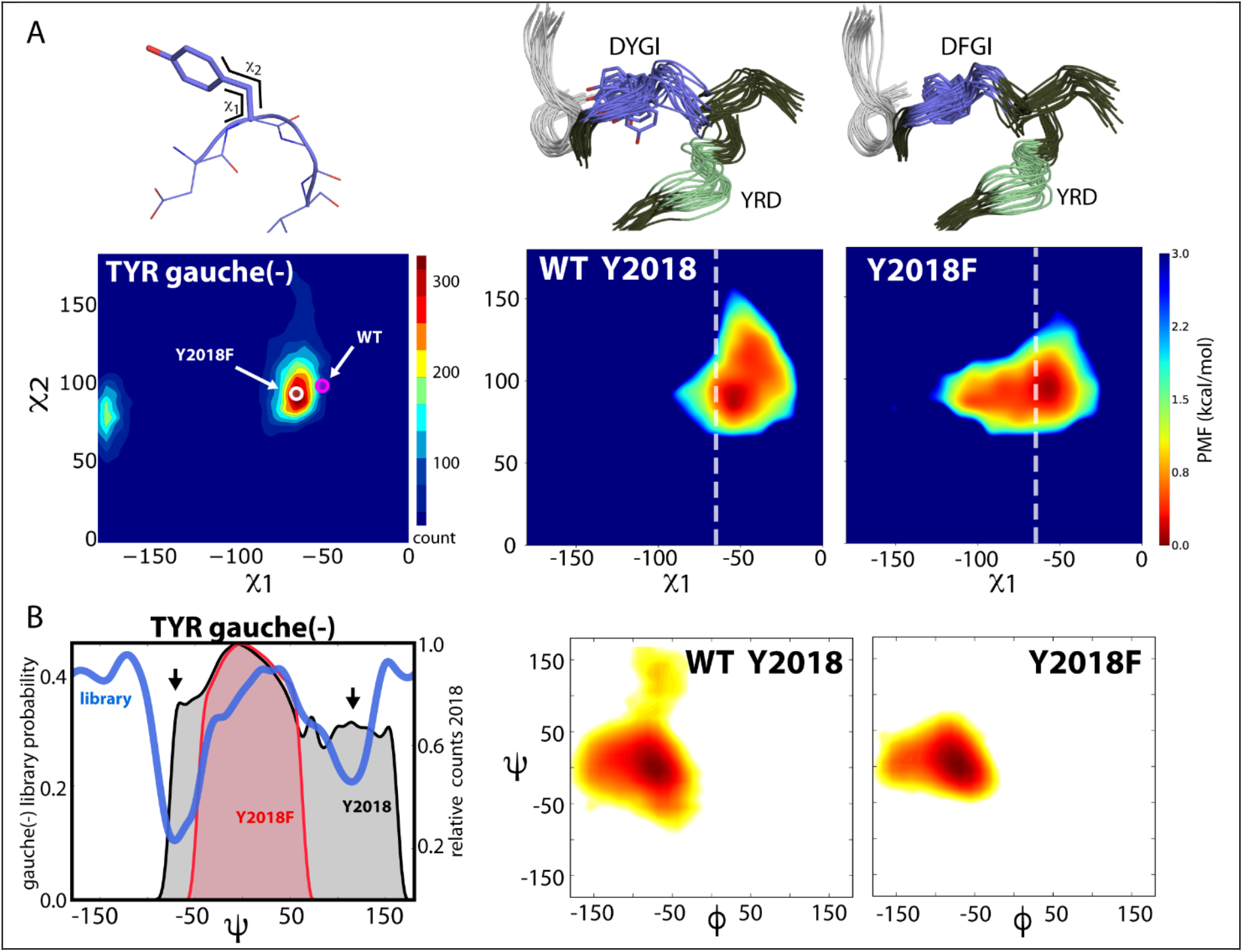
Y2018 has a non-ideal sidechain torsion angle that introduces disorder to the DYG loop. **(A)** Ideal Tyr χ1/ χ2 gauche(−) torsions from the Dunbrack database is shown with blue-to-red color indicating increasing torsion angle preference (88). The average torsion angles for wt Y2018 (magenta circle) and Y2018F (white circle) from the simulation are overlaid, showing that wt Y2018’s sidechain is in a non-preferred conformation. The χ1 / χ2 sidechain torsion angle distribution for Y2018 and Y2018F during the simulations are shown (**right panels**). Y2018F breaks the hydrogen bonds that lock the sidechain of Y2018 in wt and leads to a more ideal side-chain conformation in the mutant. The white dashed line indicates the preferred χ1 for Tyr from the Dunbrack dataset. **(B)** The ideal backbone dependent conformation for the gauche(−) rotamer of Tyr, derived from rotamer libraries (88), is depicted as the probability of the backbone ψ-dihedral angle (_i_N- _i_Cα- _i_C- _i+1_N) in blue. The non-ideal Y2018 rotamer in wt adds strain to the backbone: the Tyr backbone explores non-preferred conformations (black area, arrows indicate a non-ideal backbone ψ-angle). The backbone conformation of Y2018F populates ideal dihedral space (red area). The ϕ/ψ distribution of the backbone of Y/F2018 during the simulation is shown (**right panels**). The wt Y2018 backbone is more dynamic than Y2018F. In Y2018F the sidechain is free to access the preferred rotamer torsions and the backbone conformation is stabilized in the preferred active backbone conformation. The mutation stabilizes the entire DYGI backbone as seen in the conformational ensemble from MD (**upper right**).

**Figure S4.**
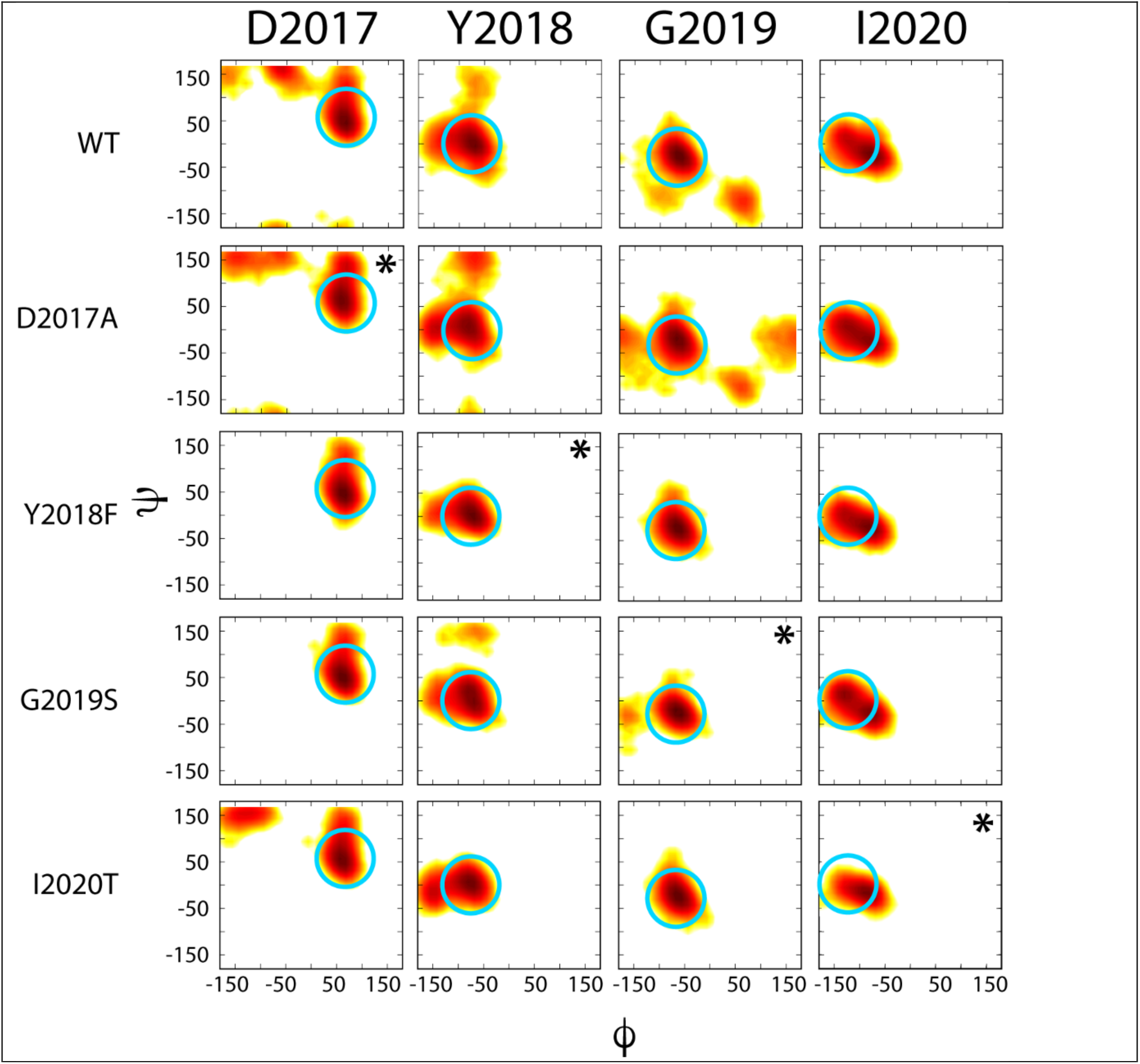
DYGI backbone dihedrals from MD simulations. DYGψ dihedral space that is associated with an activated kinase are circled in cyan (62). The wt kinase and D2017A mutant have a dynamic DYG loop that samples conformations associated with inactive kinases. The DYG loop of the mutants converge to a ϕ/ψ space that is typical of active / closed kinases. Asterisks indicate sites of mutation.

**Figure S5.**
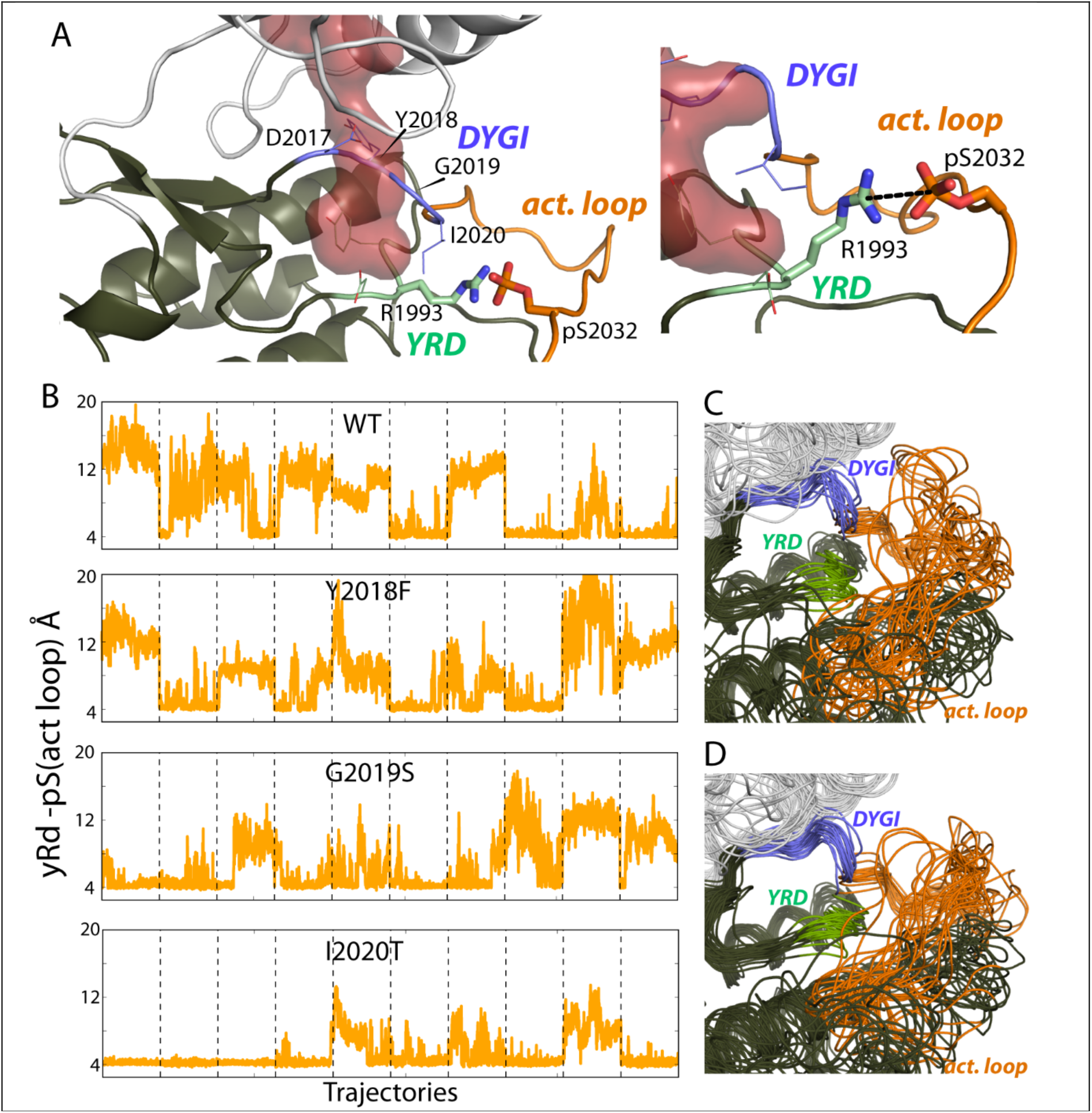
Activation loop stability from MD simulations. **(A/B)** The stability of the activation loop is represented as the distance between the YRD loop (R1993) and activation loop (pS2032). The salt-bridge between YRD and activation loop couples the catalytic loop with the N- and C-lobes. The activation loop of wt is the least stable, while activating DYGψ mutants increase stability. **(C)** The wt activation loop conformational ensemble is more widely distributed and more solvent accessible than **(D)** the I2020T activation loop. The DYGI motif is shown in blue, YRD motif in green, and activation loop in orange.

## Supplementary videos

Movie 1. Time-lapse imaging of HEK293T cells transiently expressing YFP-LRRK2-G2019S. The time interval is 5 minutes. MLi-2 was added right after the second frame. Images (640×640 pixels) of YFP fluorescence (515 nm excitation/530-630 nm emission) represent 3D volume reconstructions from confocal image stacks over time.

Movie 2. Time-lapse imaging of HEK293T cells transiently expressing YFP-LRRK2-G2019S following wash-out of MLi-2. The time interval is 11 minutes. Images (640×640 pixels) of YFP fluorescence (515 nm excitation/530-630 nm emission) represent 3D volume reconstructions from confocal image stacks over time.

Movie 3. Time-lapse imaging of HEK293T cells transiently expressing YFP-LRRK2-WT. The time interval is 5 minutes. MLi-2 was added right after the second frame. Images (640×640 pixels) of YFP fluorescence (515 nm excitation/530-630 nm emission) represent 3D volume reconstructions from confocal image stacks over time.

Movie 4. Time-lapse imaging of HEK293T cells transiently expressing YFP-LRRK2-WT following wash-out of MLi-2. The time interval is 11 minutes. Images (640×640 pixels) of YFP fluorescence (515 nm excitation/530-630 nm emission) represent 3D volume reconstructions from confocal image stacks over time.

